# An Ehrlich-inspired retrobiosynthesis of pharmaceutical scaffolds

**DOI:** 10.1101/2025.11.10.687684

**Authors:** Anastasia E.C. Rumpl, Devin Kowalski, Joshua R. Goodhew, Mika Hirano, Liam Bogucki, Carlos A. Rosa, Marc-André Lachance, Michael E. Pyne

## Abstract

Brewer’s yeast (*Saccharomyces cerevisiae*) acquires nitrogen from branched-chain and aromatic amino acids via the Ehrlich pathway, generating flavor (fusel) byproducts. Recently, diverting 4-hydroxyphenylacetaldehyde from Ehrlich catabolism of ʟ-tyrosine has enabled microbial production of opioids and other plant benzylisoquinolines. Yet, fusel metabolism is versatile in substrate scope, offering an untapped entry point for synthesizing structurally diverse aldehydes. Here, we repurpose the yeast Ehrlich pathway into a modular biocatalytic conduit for manufacturing privileged pharmaceutical alkaloids. We utilize retrobiosynthetic analysis and enzyme screening to derive scaffolds representative of solifenacin, colchicine, and ephedrine pharmaceuticals from simple amino acids. We survey wild yeasts for catabolism of ʟ-phenylglycine and demonstrate Ehrlich conversion to benzyl alcohol or (*R*)-phenylacetylcarbinol by 29 strains across nine genera. Implementing an ω-transaminase enables production of norephedrine from a simple amino acid input. This work unveils a generalizable biocatalytic route to clinically important alkaloids by exploiting metabolic logic from a yeast flavor pathway.

## INTRODUCTION

The yeast *Saccharomyces cerevisiae* acquires nitrogen from aromatic (ʟ-phenylalanine, ʟ-tryptophan, ʟ-tyrosine), branched-chain (ʟ-valine, ʟ-leucine, ʟ-isoleucine), and sulfur-containing (ʟ-methionine) amino acids via the Ehrlich pathway^1^. Catabolism begins with the transamination of an amino acid into its corresponding ⍺-keto acid, which is subsequently decarboxylated to an aldehyde. Depending on cultivation conditions, aldehydes are reduced to fusel alcohols or oxidized to fusel acids^2^. Derivative fusel alcohols can be further converted to acetate esters^3^. Fusel alcohols and esters are paramount as organoleptics in fermented beverages produced by *S. cerevisiae* and other brewing yeasts^4^. In recent years, microbial cell factories have been engineered for scalable synthesis of fusel products, given their applications as aromatics, solvents, and biofuels^5–8^. Reconstruction of the Ehrlich pathway in *Escherichia coli* has enabled production of the biofuel isobutanol at titers up to 50 g/L^9^, while cell-free isobutanol production has reached 275 g/L and near-theoretical yield (95%)^10^.

More recently, rewiring of the Ehrlich pathway in *S. cerevisiae* has enabled sustainable synthesis of high-value plant natural products. The central focus of these efforts has been the benzylisoquinoline alkaloids (BIAs), a class of approximately 2,500 plant secondary metabolites, many of which exhibit potent pharmacological properties^11–13^. Yeast BIA biosynthesis is possible because a key pathway precursor, 4-hydroxyphenylacetaldehyde (4-HPAA), arises from Ehrlich catabolism of ʟ-tyrosine^14^. In plants, the committed step in BIA biosynthesis involves a Pictet– Spengler condensation of 4-HPAA and dopamine to yield (*S*)-norcoclaurine^15^. Four subsequent enzymatic steps convert (*S*)-norcoclaurine to (*S*)-reticuline, a key precursor to many downstream BIAs^16^. Near-commercial production of (*S*)-reticuline has been achieved in yeast at titers up to 4.8 g/L^17^, underscoring the potential of the Ehrlich pathway as a conduit for manufacturing pharmaceutical scaffolds^18^. The (*S*)-reticuline pathway has been extended to produce numerous functionalized BIA end products, including the antitussive noscapine, the antimicrobial sanguinarine, the antispasmodic papaverine, and the opioid analgesics morphine, codeine, and hydrocodone^17,19–24^.

Condensation of 4-HPAA and dopamine is catalyzed by (*S*)-norcoclaurine synthase (NCS), a functionally diverse enzyme class^25,26^. Natural and engineered NCS variants accept a wide range of carbonyl substances, enabling the synthesis of remarkably diverse substituted tetrahydroisoquinolines (THIQs)^27,28^. Recently, by coupling dopamine biosynthesis and NCS from Japanese goldthread (*Coptis japonica*; *Cj*NCS) to the yeast Ehrlich pathway, a plethora of new-to-nature substituted THIQs were synthesized in *S. cerevisiae*^18^. Substituted THIQs were first synthesized from endogenous fusel aldehydes produced by catabolism of canonical Ehrlich pathway amino acids (ʟ-phenylalanine, ʟ-tryptophan, ʟ-tyrosine, ʟ-leucine, and ʟ-methionine). This approach was further extended by growing engineered strains on non-standard amino acids, such as ʟ-norvaline, ʟ-norleucine, and ʟ-2-aminoheptanoic acid, illustrating the broad substrate specificity of both *Cj*NCS and the Ehrlich pathway. Such extensive substrate promiscuity positions the Ehrlich pathway as an untapped tool for aldehyde production and potentially illuminates new biocatalytic routes to broad aldehyde-based pharmaceuticals.

Here, we design, construct, and validate a generalizable retrobiosynthetic route to several classes of approved pharmaceutical compounds. Leveraging the yeast Ehrlich pathway as an entry point to structurally diverse aldehydes, we retrobiosynthetically predict and experimentally validate production of alkaloid scaffolds involved in the synthesis of the pharmaceuticals solifenacin (overactive bladder), colchicine (gout), and ephedrine (hypotension). We show improved production of alkaloid scaffolds by blocking the conversion of aldehyde precursors to fusel alcohols and screening wild yeast strains for improved amino acid catabolism. To showcase the broader utility of our approach, we demonstrate conversion of an alkaloid scaffold [(*R*)-phenylacetylcarbinol] to an active pharmaceutical ingredient (norephedrine) starting from a simple input (ʟ-phenylglycine). Collectively, our retrobiosynthetic approach unveils bespoke pathways to clinically important alkaloids through rewiring of yeast amino acid catabolism.

## RESULTS

### Screening diverse NCS orthologs for production of THIQs from endogenous amino acids

Previously, it was shown that *S. cerevisiae* strains harboring a dopamine biosynthesis pathway and a truncated NCS from *C. japonica* (*Cj*NCSΔN) convert endogenous Ehrlich pathway amino acids (ʟ-Tyr, ʟ-Phe, ʟ-Trp, ʟ-Leu, and ʟ-Met) to the corresponding aldehydes, which condense with dopamine to give an array of substituted THIQs^18^ (**Fig. 1a**). Based on this general pathway, we wished to construct a library of NCS variants to screen for conversion of broad Ehrlich pathway aldehydes to substituted THIQs. Using literature and homology searches, we selected 10 plant NCS variants to screen, which included C- and N-terminal truncation variants reported in the literature (e.g., *Sd*NCSΔC from *Stylophorum diphyllum* and *Sc*NCSΔN from *Sanguinaria canadensis*)^29^ and engineered mutants with altered substrate specificities (*Tf*NCS-M97V from *Thalictrum flavum*)^30^. We also included *Cj*NCSΔN, which has been widely utilized for high-level BIA production in yeast^17,18^ (**Supplementary Table 1; Supplementary Figure 1**).

**Figure 1.**
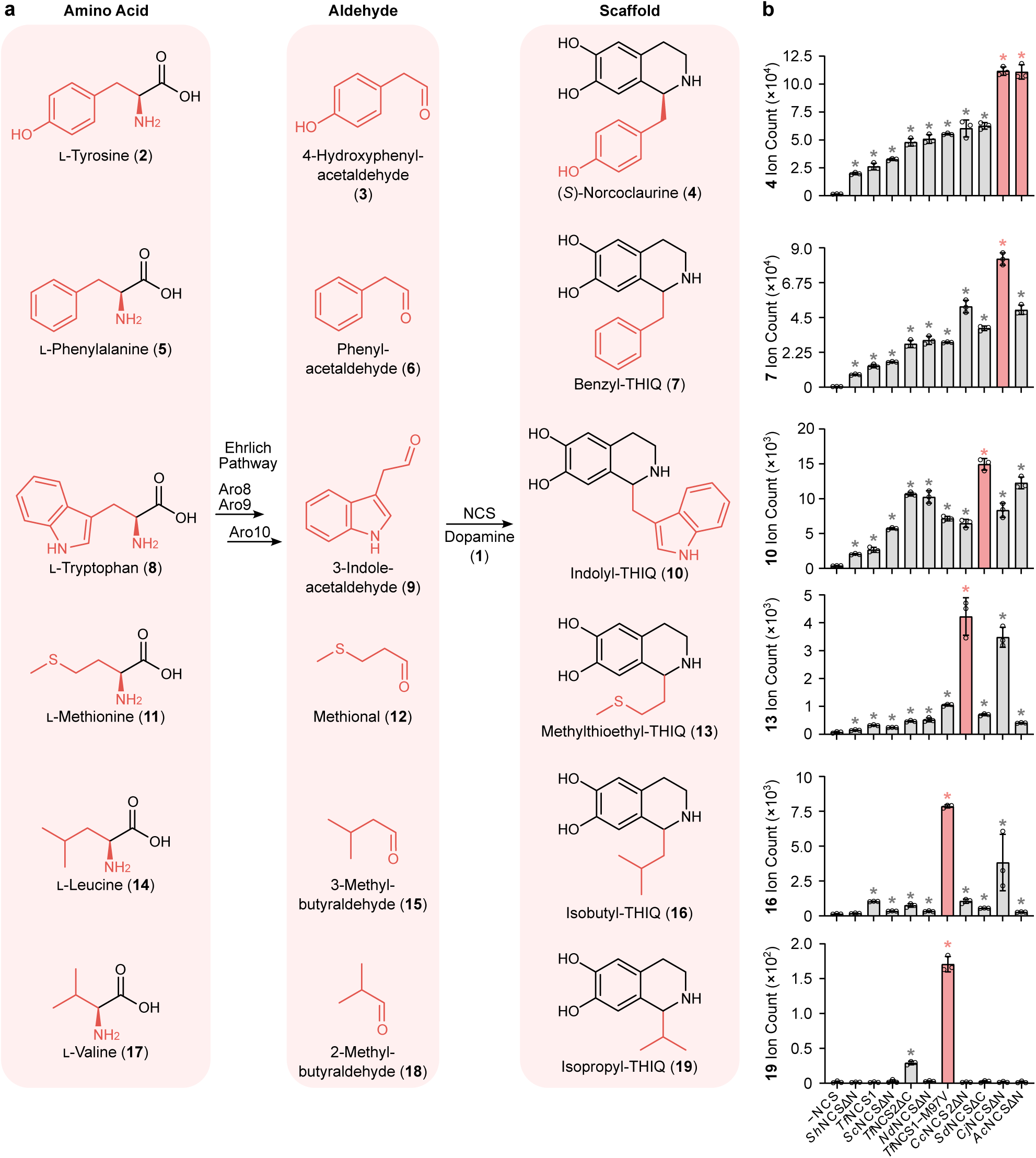
Screening a library of norcoclaurine synthase (NCS) variants for production of substituted tetrahydroisoquinolines (THIQs) from endogenous amino acids. **a,** Metabolic pathways for production of substituted THIQs by redirecting fusel aldehydes from the Ehrlich pathway (Aro8/Aro9 and Aro10). Incorporation of amino acids into aldehydes and substituted THIQs is shown in red. Stereochemistry is shown for (*S*)-norcoclaurine and omitted from other substituted THIQs. b, Plant NCS variants differ in their ability to convert aliphatic and aromatic aldehydes into substituted THIQs. Engineered yeast strains were grown on a mixture of Ehrlich pathway amino acids [ʟ-Tyr (2), ʟ-Phe (5), ʟ-Trp (8), ʟ-Met (11), ʟ-Leu (14), and ʟ-Val (17)] as a source of nitrogen to promote formation of substituted THIQs. NCS variants leading to formation of the greatest amount of target THIQ product are shown in red. Asterisk (*) denotes a significant increase (*P* < 0.05) in target ion count relative to the control strain lacking an NCS enzyme. Error bars represent the mean ± s.d. of *n* = 3 independent biological replicates. Statistical differences between control and derivative strains were tested using two-tailed Welch’s *t*-test. Abbreviations: NCS, norcoclaurine synthase; THIQ, tetrahydroisoquinoline.

NCS-encoding genes were chromosomally integrated into a dopamine (**1**) production host (*PpDODC* from *Pseudomonas putida* and *BvCYP76AD5* from *Beta vulgaris*^16^) possessing a deregulated and overexpressed ʟ-tyrosine (**2**) pathway (*ARO4^K229L^*, *ARO7^G141S^*, and *TYR1*). NCS variant strains were transformed with pHLUM plasmid containing *HIS3*, *LEU2*, *URA3*, and *MET17* markers to complement auxotrophic deletions and restore prototrophy^31^. Prototrophic NCS strains were grown in 2× YNB medium containing a mixture of Ehrlich pathway amino acids [ʟ-Tyr (**2**), ʟ-Phe (**5**), ʟ-Trp (**8**), ʟ-Met (**11**), ʟ-Leu (**14**), and ʟ-Val (**17**)] as a source of nitrogen. Cultures were assayed for the presence of corresponding substituted THIQs, which revealed differences in the substrate scope of engineered NCS variants (**Fig. 1b**). *Ac*NCSΔN, a previously uncharacterized enzyme from blue columbine (*Aquilegia coerulea*), yielded comparable levels of (*S*)-norcoclaurine (**4**) relative to *Cj*NCSΔN, the most efficient NCS variant reported to date^17,18^. However, *Ac*NCSΔN exhibited greater selectivity for (*S*)-norcoclaurine (**4**) compared to *Cj*NCSΔN, as *Ac*NCSΔN did not efficiently convert aldehydes derived from ʟ-Met [methional (**12**)] or ʟ-Leu [(3-methylbutyraldehyde (**15**)]. *Sd*NCSΔC and *Tf*NCS1-M97V exhibited the greatest activity on aldehydes derived from bulky (ʟ-Trp) and branched-chain (ʟ-Leu) amino acids, respectively. In agreement with its altered substrate scope^30^, *Tf*NCS1-M97V accepted 2-methylbutyraldehyde (**18**), a challenging α-substituted aldehyde derived from ʟ-Val catabolism^1^. Yeast-derived THIQ products co-eluted with and generated identical fragmentation spectra to synthetic THIQ standards prepared through the phosphate-catalyzed condensation of Ehrlich pathway aldehydes with dopamine (**Supplementary Figure 2**). In agreement with prior studies^16,18^, chiral analysis of norcoclaurine revealed that all NCS variants stereospecifically produced (*S*)-norcoclaurine relative to a racemic standard (**Supplementary Figure 3**). All NCS variants also yielded enantiopure benzyl-THIQ derived from ʟ-Phe, yet absolute stereochemistry could not be assigned due to the absence of an enantiopure standard.

### Retrobiosynthesis of pharmaceutical tetrahydroisoquinolines from non-standard amino acids

The capacity of *S. cerevisiae* to derive nitrogen from structurally diverse amino acids^1^, coupled with the tremendous substrate scope of plant NCS enzymes^28^, led us to speculate that other families of natural and medicinal THIQs could be accessed using the Ehrlich pathway. In nature, plants synthesize more than 2,000 BIAs (benzyl-THIQs) from ʟ-Tyr via the C_6_-C_2_ aldehyde 4-hydroxyphenylacetaldehyde [4-HPAA (**3**)]^11^. Beyond BIAs, other classes of THIQs have been identified in plants. Autumn crocus (*Colchicum autumnale*) and flame lily (*Gloriosa superba*) synthesize the phenethylisoquinoline (phenethyl-THIQ) drug colchicine (**28**) from 4-hydroxydihydrocinnamaldehyde [4-HDCA (**26**)], a C_6_-C_3_ aldehyde derived from the phenylpropanoid pathway^32^. Further, species of orchids from the genus *Cryptostylis* synthesize phenylisoquinolines (phenyl-THIQs), such as cryptostyline I, II, and III, presumably from C_6_-C_1_ benzaldehydes^33,34^. Solifenacin is a synthetic phenyl-THIQ used to treat overactive bladder and other phenyl-THIQs have shown promising activities against cancer and HIV^35,36^.

Based on retrosynthetic analysis using the Ehrlich pathway, phenyl-THIQs might be accessible through yeast catabolism of ʟ-phenylglycine [ʟ-Phg (**20**)] via the C_6_-C_1_ intermediate benzaldehyde (**21**), while catabolism of ʟ-homotyrosine [ʟ-Hty (**25**)] is expected to yield 4-hydroxyphenethyl-THIQs from the C_6_-C_3_ aldehyde 4-HDCA (**26**) (**Fig. 2a**). To probe these theoretical routes, we first attempted to grow prototrophic *S. cerevisiae* BY4741 on ʟ-Phg (**20**) or ʟ-Hty (**25**) as sole nitrogen sources, as growth on these ʟ-Phe (**5**) and ʟ-Tyr (**2**) analogs would imply catabolism via the Ehrlich pathway. Relative to a control lacking nitrogen (Area Under the Curve, AUC = 51.0 ± 1.0), *S. cerevisiae* was able to grow on ʟ-Phg (AUC = 79.0 ± 1.0) or ʟ-Hty (AUC = 87.0 ± 2.0) as sole nitrogen sources, albeit to a lesser extent than growth on ʟ-Phe (AUC = 110.7 ± 1.2) and ʟ-Tyr (AUC = 99.3 ± 1.5) (**Supplementary Figure 4**).

**Figure 2.**
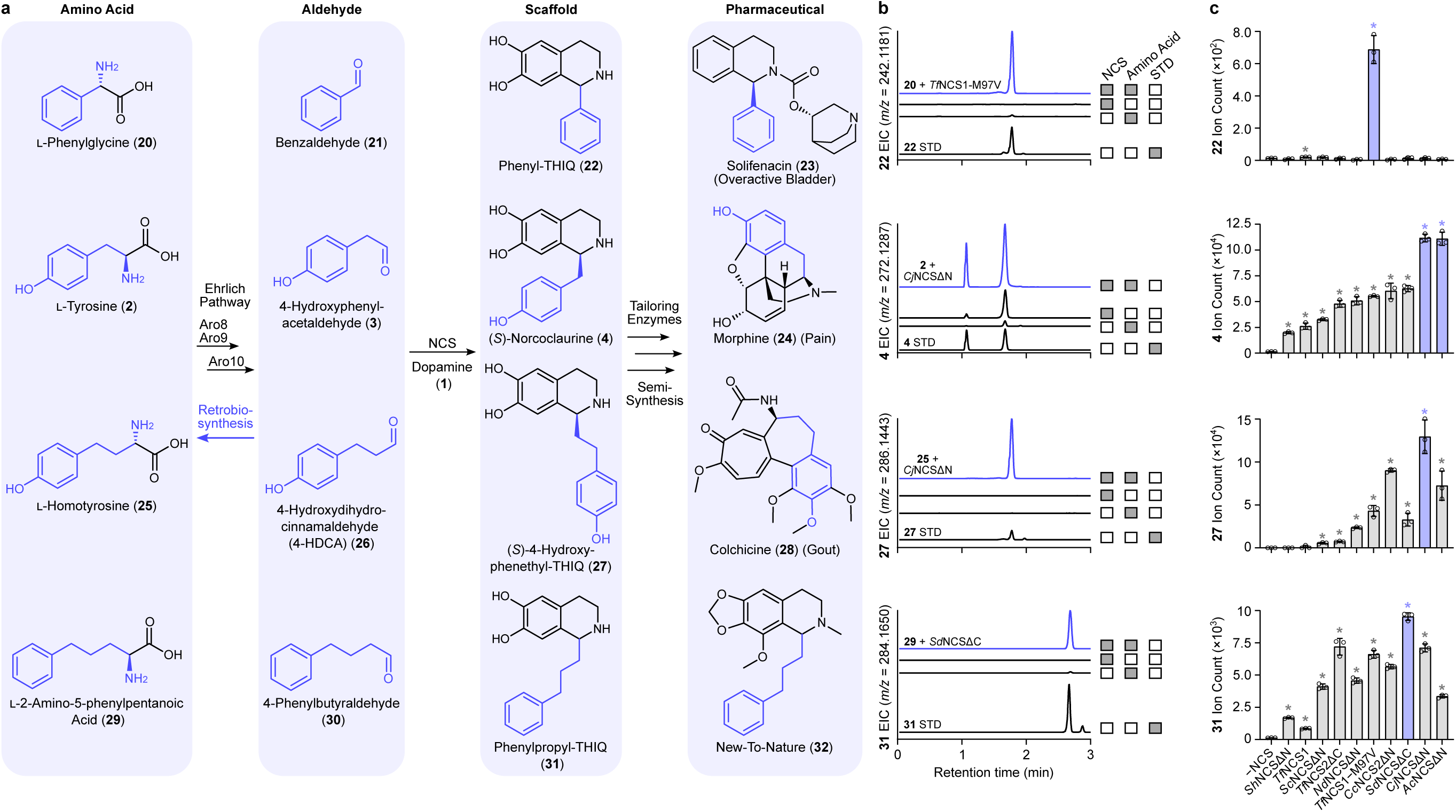
Retrobiosynthesis of pharmaceutical tetrahydroisoquinolines (THIQs) from non-standard amino acids. **a,** Proposed retrobiosynthetic pathways for production of pharmaceutical THIQs from non-standard amino acids via the Ehrlich pathway (Aro8/Aro9 and Aro10). THIQ scaffolds are converted into approved pharmaceuticals or new-to-nature structures (**32**) by plant tailoring enzymes or chemical methods (semi-synthesis). Incorporation of non-standard amino acids into aldehydes, substituted THIQs, and pharmaceuticals or new-to-nature compounds is shown in blue. Conversion of ʟ-tyrosine into morphine is shown for comparison. Proposed retrobiosynthetic routes from aldehydes to non-standard amino acids is shown with a blue arrow. Stereochemistry is shown for (*S*)-norcoclaurine and (*S*)-4-hydroxyphenethyl-THIQ; other THIQs are produced stereospecifically, yet absolute stereochemistry could not be determined. **b,** Ion-extracted LC-MS chromatograms of yeast strains harboring a dopamine biosynthesis pathway and a plant NCS variant grown on various non-standard amino acids as a sole nitrogen source. Retention times of target THIQ scaffolds are compared against authentic THIQ standards prepared through the *in vitro* condensation of dopamine and the corresponding aldehyde. Control chromatograms derived from strains lacking an NCS enzyme or supplemented with ammonium sulfate in place of non-standard amino acids are shown for comparison. NCS-derived (*S*)-norcoclaurine and commercial (*R,S*)-norcoclaurine generate two LC-MS peaks. LC-MS chromatographic experiments were performed three times with each replicate yielding similar results. **c,** Screening a library of norcoclaurine synthase (NCS) variants enables production of pharmaceutical THIQ scaffolds. Engineered yeast strains were grown on non-standard amino acids [ʟ-Phg (**20**), ʟ-Hty (**25**), or ʟ-2-amino-5-phenylpentanoic acid (ʟ-APPA, **29**)] as a sole source of nitrogen to promote formation of substituted THIQs (**22**, **27**, **31**). NCS variants leading to formation of the greatest amount of target THIQ product are shown in blue. Asterisk (*) denotes a significant increase (*P* < 0.05) in target THIQ product relative to the control strain lacking an NCS enzyme. Error bars represent the mean ± s.d. of *n* = 3 independent biological replicates. Statistical differences between control and derivative strains were tested using two-tailed Welch’s *t*-test. Abbreviations: EIC, extracted ion chromatogram; min, minute; NCS, norcoclaurine synthase; STD, standard; THIQ, tetrahydroisoquinoline.

To determine if catabolism of ʟ-Phg and ʟ-Hty proceeds via the proposed Ehrlich route and results in formation of the corresponding aldehydes (benzaldehyde and 4-HDCA, respectively) (**Fig. 2a**), we cultivated 10 NCS-containing yeast strains on ʟ-Phg or ʟ-Hty and assayed cultures for the presence of the respective phenyl-THIQ (**22**) and 4-hydroxyphenethyl-THIQ (**27**) scaffolds (**Fig. 2b**). Following growth of our NCS strain library on ʟ-Phg as sole nitrogen source, only two strains (*Tf*NCS1 and *Tf*NCS1-M97V) yielded LC-MS peaks consistent with the expected phenyl-THIQ product (*m/z* 242.1181 ± 7.0 ppm) (**Fig. 2c**). Yeast-derived phenyl-THIQ yielded an identical retention time and fragmentation spectra in agreement with phenyl-THIQ standard prepared through *in vitro* condensation of benzaldehyde (**21**) and dopamine (**1**) (**Fig.** 2b, Supplementary Figure 4).

Analogously, following growth of our NCS strain library on ʟ-Hty, several NCS strains synthesized 4-hydroxyphenethyl-THIQ (**27**) consistent with condensation of dopamine (**1**) and 4-HDCA (**26**) (*m/z* 286.1443 ± 1.0 ppm) (**Fig. 2c**). Strains containing *Cj*NCSΔN produced the greatest amount of 4-hydroxyphenethyl-THIQ, followed by *Cc*NCS2ΔN and *Ac*NCSΔN. Yeast-derived 4-hydroxyphenethyl-THIQ co-eluted with and yielded an identical fragmentation spectra to a 4-hydroxyphenethyl-THIQ standard derived through *in vitro* condensation of 4-HDCA and dopamine (**Fig. 2b**, **Supplementary Figure 4**). Similarly, growth of our NCS strain library on ʟ-homophenylalanine [ʟ-Hph (**33**)] yielded a phenethylisoquinoline (phenethyl-THIQ) product (**35**) (*m/z* 270.1494 ± 5.9 ppm) with an identical retention time and fragmentation spectra as a phenethyl-THIQ standard generated from the *in vitro* condensation of 3-phenylpropionaldehyde (**34**) and dopamine (**1**) (**Supplementary Figure 5**).

Having demonstrated synthesis of phenyl-THIQ and phenethyl-THIQ scaffolds from respective C_6_-C_1_ and C_6_-C_3_ aldehydes, we wished to extend our retrobiosynthetic approach to the phenylpropylisoquinolines (phenylpropyl-THIQ). Phenylpropyl-THIQs are not found in nature, yet a synthetic phenylpropyl-THIQ (**32**) has been reported to exhibit analgesic properties^37^. Using our Ehrlich-inspired retrobiosynthesis pathway, the phenylpropyl-THIQ scaffold (**31**) can be accessed through catabolism of ʟ-2-amino-5-phenylpentanoic acid [ʟ-APPA (**29**)] via the C_6_-C_4_ aldehyde 4-phenylbutyraldehyde (**30**). Prototrophic *S. cerevisiae* BY4741 was able to utilize ʟ-APPA as a sole nitrogen source (AUC = 83.3 ± 0.6) to a similar extent as growth on ʟ-Phg (AUC = 79.0 ± 1.0) and ʟ-Hty (AUC = 87.0 ± 2.0). Growth of our NCS strain library on ʟ-APPA as a sole source of nitrogen revealed several NCS variants with the ability to produce phenylpropyl-THIQ (**32**) (*m/z* 284.1650 ± 1.8 ppm) (**Fig. 2c)**. *Sd*NCSΔC yielded the greatest amount of phenylpropyl-THIQ (**32**), consistent with its ability to accept the bulky 3-indole acetaldehyde (**9**) substrate derived from Ehrlich catabolism of ʟ-Trp (**Fig. 1b)**. Yeast-derived phenylpropyl-THIQ co-eluted with and yielded an identical fragmentation spectra to a standard derived through *in vitro* condensation of 4-phenylbutyraldehyde and dopamine (**Fig. 2b**, **Supplementary Figure 4**).

In all experiments involving non-standard amino acids (ʟ-Phg, ʟ-Hph, ʟ-Hty, or ʟ-APPA), omitting an NCS biosynthetic enzyme or replacing the non-standard amino acid with ammonium sulfate as the source of nitrogen failed to yield the expected substituted THIQ products (**Fig. 2b, Supplementary Figure 5**), confirming that THIQ products arise from Ehrlich catabolism of non-standard amino acids. Chiral analysis of THIQs derived from non-standard amino acids revealed that all active NCS variants produced enantiopure THIQ products relative to racemic standards (**Supplementary Figure 6**). Absolute stereochemistry of THIQ products could not be assigned due to the absence of enantiopure standards, except for (*S*)-4-hydroxyphenethyl-THIQ (**27**) derived from ʟ-Hty, as *Cj*NCSΔN has been shown to stereospecifically produce (*S*)-4-hydroxyphenethyl-THIQ from dopamine and 4-HDCA^32^.

### Retrobiosynthesis of substituted amphetamines from ʟ-phenylglycine

After successfully synthesizing diverse substituted THIQ scaffolds from C_6_-C_1_, C_6_-C_3_, and C_6_-C_4_ aldehydes, we sought to extend our retrobiosynthetic strategy to another alkaloid class—the substituted amphetamines. The key substituted amphetamine scaffold, (*R*)-phenylacetylcarbinol (PAC), has been used in the chemical synthesis of ephedrine for over a century^38^. PAC is synthesized by yeast via condensation of pyruvate with supplemented benzaldehyde by endogenous pyruvate decarboxylase (PDC). Having demonstrated that Ehrlich catabolism of ʟ-Phg generates phenyl-THIQ via benzaldehyde (**Fig. 2**), we reasoned that ʟ-Phg could also serve as a precursor to PAC (**Fig. 3a**). Accordingly, *S. cerevisiae* BY4741 was transformed with pHLUM to restore prototrophy and grown on ʟ-Phg as a sole source of nitrogen. Under these conditions, HPLC-MS confirmed detection of PAC in culture extracts, verifying the capacity of *S. cerevisiae* to convert ʟ-Phg to PAC (*m/z* – H_2_O = 133.0653 ± 1.0 ppm) (**Fig. 3b**). Yeast-derived PAC co-eluted with and generated identical fragmentation spectra as an authentic PAC standard (**Fig. 3c**). Substitution of ʟ-Phg with ammonium sulfate as the sole nitrogen source abolished PAC production, confirming that PAC originates from ʟ-Phg catabolism.

**Figure 3.**
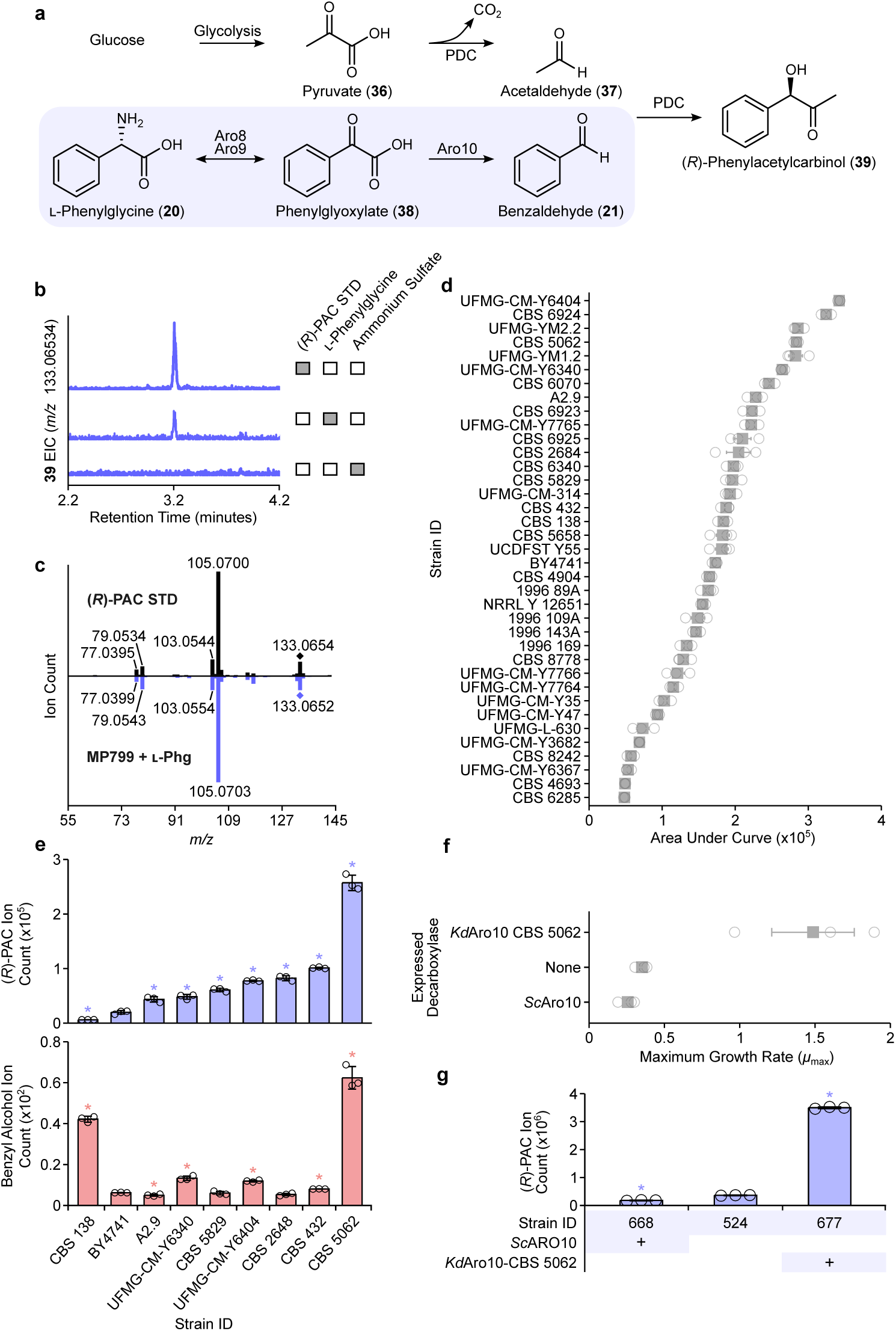
Retrobiosynthesis of substituted amphetamines from ʟ-phenylglycine (ʟ-Phg). **a,** Proposed retrobiosynthetic pathway for production of (*R*)-phenylacetylcarbinol [(*R*)-PAC] via Ehrlich catabolism of ʟ-Phg (Aro8/Aro9 and Aro10; shaded). Condensation of benzaldehyde with acetaldehyde by endogenous pyruvate decarboxylases (PDCs) yields (*R*)-PAC. **b,** Ion-extracted LC-MS chromatogram of *S. cerevisiae* BY4741 grown on ʟ-Phg as a sole nitrogen source. Retention time of yeast-produced (*R*)-PAC is compared against an authentic commercial standard. A control chromatogram derived from *S. cerevisiae* BY4741 supplemented with ammonium sulfate in place of ʟ-Phg is shown. **c,** MS/MS fragmentation spectra of commercial (*R*)-PAC standard and spent media from *S. cerevisiae* strain MP799 grown on ʟ-Phg as a sole nitrogen source. **d,** Growth of non-conventional yeasts and an *S. cerevisiae* BY4741 control grown on ʟ-Phg as a sole nitrogen source. Growth was assessed using Area Under the Curve (AUC). **e,** End product metabolites [PAC (blue) and benzyl alcohol (pink)] synthesized by *S. cerevisiae* BY4741 and select non-conventional yeasts grown on ʟ-Phg as a sole nitrogen source. **f,** Maximum specific growth rate (*µ*_max_) of engineered *S. cerevisiae* strains expressing heterologous *KdARO10* from *K. dobzhanskii* CBS 5062 or overexpressing native *ScARO10* from *S. cerevisiae* BY4741. Engineered strains were grown on ʟ-Phg as a sole nitrogen source and compared against a control strain lacking *ARO10* overexpression. **g,** PAC production by engineered *S. cerevisiae* strains expressing *KdARO10* or overexpressing *ScARO10*. Engineered strains were grown on ʟ-Phg as a sole nitrogen source and compared against a control strain lacking *ARO10* overexpression. Asterisk (*) denotes a significant increase or decrease (*P* < 0.05) in PAC relative to a control strain lacking an overexpressed *ARO10*. Error bars represent the mean ± s.d. of *n* = 3 independent biological samples. Statistical differences were tested using two-tailed Welch’s *t*-test. LC-MS chromatographic and MS/MS fragmentation experiments were performed three times and two times, respectively, with each replicate yielding similar results. Abbreviations: AUC, area under the curve; EIC, extracted ion chromatogram; (*R*)-PAC, (*R*)-phenylacetylcarbinol; PDC, pyruvate decarboxylase; ʟ-Phg, ʟ-phenylglycine; STD, standard.

Genomic variation among yeasts can lead to differences in substrate catabolism, including the utilization of nitrogen sources^39^. Despite requiring no additional heterologous enzymes, ʟ-Phg conversion to PAC by *S. cerevisiae* BY4741 was notably weak. Given that fusel metabolism has been observed across multiple orders within the Ascomycota and even among members of the Basidiomycota^14^, we reasoned that alternative non-*Saccharomyces* yeasts may exhibit enhanced Ehrlich catabolism of ʟ-Phg and conversion to PAC. Accordingly, we selected and screened 36 non-conventional yeast strains from 11 different genera for their ability to grow on ʟ-Phg as a sole nitrogen source (**Supplementary Table 2**). Overall, most yeasts exhibited measurable growth relative to an *S. cerevisiae* control lacking nitrogen, with 19 strains outperforming *S. cerevisiae* BY4741 (**Fig. 3d**). The five best-growing strains exhibited AUCs 1.6- to 2-fold higher than that of *S. cerevisiae* BY4741 (AUC = 17.3 × 10^5^ ± 0.4 × 10^5^), including *Zygotorulaspora cariocana* UFMG-CM-Y6404 (AUC = 34.2 × 10^5^ ± 0.2 × 10^5^), *Lachancea* sp. CBS 6924 (AUC = 32.4 × 10^5^ ± 0.7 × 10^5^), *Lachancea nothofagi* UFMG-YM2.2 (AUC = 28.6 × 10^5^ ± 0.8 × 10^5^), *Kluyveromyces dobzhanskii* CBS 5062 (AUC = 28.4 × 10^5^ ± 0.3 × 10^5^), and *Lachancea nothofagi* UFMG-YM1.2 (AUC = 28.3 × 10^5^ ± 1.6 × 10^5^). We subsequently selected 30 strains with the highest AUCs for HPLC-MS analysis of ʟ-Phg-derived end products (PAC and benzyl alcohol). All 30 yeasts produced benzyl alcohol, while nine produced PAC (**Supplementary Table 3**). *K. dobzhanskii* CBS 5062 generated the highest concentrations of ʟ-Phg-derived metabolites, synthesizing 13-fold more PAC and 10-fold more benzyl alcohol than *S. cerevisiae* BY4741.

To improve PAC production by *S. cerevisiae*, we focused on the irreversible conversion of the ⍺-keto acid phenylglyoxylate to benzaldehyde by the presumed phenylpyruvate decarboxylase Aro10. The Aro10-encoding gene from *Kluyveromyces dobzhanskii* CBS 5062 (*KdARO10*) and *S. cerevisiae* BY4741 (*ScARO10*) were chromosomally integrated into a PAC production host containing *Ct*PDC from *Candida tropicalis*. The Aro10 variant strains were subsequently transformed with pHLUM to complement auxotrophic deletions and grown in medium containing ʟ-Phg as a sole source of nitrogen. Relative to the parental host (*µ*_max_ = 0.35 ± 0.04), the strain expressing heterologous *KdARO10* (*µ*_max_ = 1.49 ± 0.47) demonstrated improved growth on ʟ-Phg (**Fig. 3e**). Cultures were assayed for the presence of PAC, which revealed differences in the substrate scope of Aro10 variants. While expression of heterologous *KdARO10* improved PAC production 10-fold, overexpression of *ScARO10* decreased PAC titer 2-fold (**Fig. 3f**).

### Deleting host oxidoreductases to improve production of alkaloid scaffolds

Microbial strains engineered to produce aldehyde-derived metabolites are limited by the redox conversion of aldehydes to unwanted side products by redundant host oxidoreductases^18,40,41^. Accordingly, we reasoned that the synthesis of substituted THIQ and amphetamine scaffolds by our engineered strains was limited by the conversion of Ehrlich-derived aldehyde precursors (benzaldehyde, 4-HDCA, and 4-phenylbutyraldehyde) to corresponding alcohol side products. Notably, the reduction of benzaldehyde to benzyl alcohol has presented as a century-long obstacle to efficient PAC synthesis in yeast, yet little progress has been made in identifying the culprit oxidoreductases^42,43^. To improve aldehyde availability, we selected nine NADPH-dependent *S. cerevisiae* oxidoreductases (Adh6, Ari1, Gcy1, Gre2, Gre3, Hfd1, Ypr1, Ydr541c, Ygl039w), for deletion screening in each scaffold-producing strain, based on their involvement in the reduction of broad aldehydes to fusel alcohols within the Ehrlich pathway^14,40,44^.

Repeating the oxidoreductase deletion screen in our substituted amphetamine platform strain implicated Adh6, Ari1, Gcy1, Gre2, Gre3, and Ypr1 in benzaldehyde reduction, as their inactivation facilitated significant improvements in (*R*)-PAC production (**Fig. 4a**). To further shut off benzaldehyde reduction, five of the aldo-keto reductases (Adh6, Ari1, Gcy1, Gre2, Ypr1) were concurrently knocked out of our host, enabling a 3.2-fold decrease in benzyl alcohol production (**Fig. 4b**). Overall, combined inactivation of the five oxidoreductases increased PAC synthesis 2.2-fold (**Fig. 4c**).

**Figure 4.**
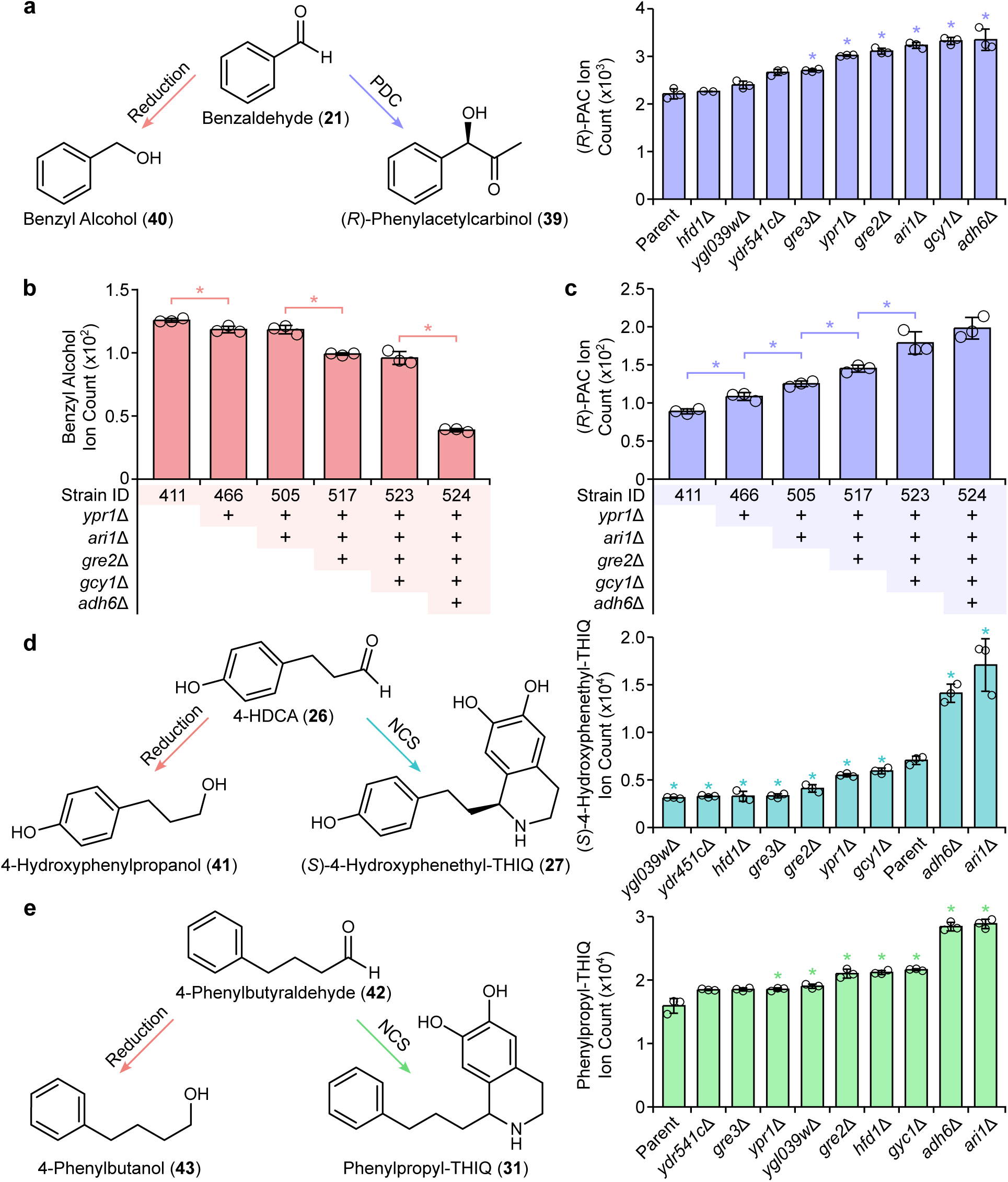
Improving pharmaceutical scaffold production via host oxidoreductase deletions. **a,** Benzaldehyde is reduced to benzyl alcohol by unidentified yeast oxidoreductases in industrial production of (*R*)-phenylacetylcarbinol (PAC). Yeast strains with single NADPH-dependent oxidoreductase deletions were grown on benzaldehyde alongside a parental control. All strains harbor a pyruvate decarboxylase (PDC) variant from *Candida tropicalis* (*Ct*PDC) **b,** Benzyl alcohol formation from exogenous benzaldehyde by yeast strains containing up to five aldo-keto reductase deletions. **c,** (*R*)-PAC formation from exogenous benzaldehyde by yeast strains containing up to five aldo-keto reductase deletions. **d,** Production of (*S*)-4-hydroxyphenethyl-THIQ from ʟ-homotyrosine by yeast strains harboring single oxidoreductase gene deletions. ʟ-Homotyrosine (**25**) is converted to 4-HDCA (**26**) and reduced by host oxidoreductases to the corresponding alcohol [4-hydroxyphenylpropanol (**41**)] or condensed with dopamine by NCS to form (*S*)-4-hydroxyphenethyl-THIQ (**27**). All strains possess a dopamine biosynthesis pathway and *Ac*NCSΔN from *Aquilegia coerulea*. **e,** Production of phenylpropyl-THIQ from ʟ-2-amino-5-phenylpentanoic acid (ʟ-APPA) by yeast strains harboring single oxidoreductase gene deletions. ʟ-APPA (**29**) is converted to 4-phenylbutyraldehyde (**30**) and reduced by host oxidoreductases to the corresponding alcohol [4-phenylbutanol (**43**)] or condensed with dopamine by NCS to form phenylpropyl-THIQ (**31**). All strains possess a dopamine biosynthesis pathway and *Ac*NCSΔN from *Aquilegia coerulea*. Asterisk (*) denotes a significant increase or decrease (*P* < 0.05) in (*R*)-PAC, benzyl alcohol, (*S*)-4-hydroxyphenethyl-THIQ, or phenylpropyl-THIQ relative to parent strains. Error bars represent the mean ± s.d. of *n* = 3 independent biological samples. Statistical differences between control and derivative strains were tested using two-tailed Welch’s *t*-test. Abbreviations: 4-HDCA, 4-hydroxydihydrocinnamaldehyde; NCS, norcoclaurine synthase; (*R*)-PAC, (*R*)-phenylacetylcarbinol; PDC, pyruvate decarboxylase; THIQ, tetrahydroisoquinoline.

We next sought to investigate the effect of inactivating the same nine NADPH-dependent oxidoreductases (Adh6, Ari1, Gcy1, Gre2, Gre3, Hfd1, Ypr1, Ydr541c, Ygl039w) on production of 4-hydroxyphenethyl-THIQ (**27**) and phenylpropyl-THIQ (**31**) scaffolds. Deletion screening yielded marked improvements in the synthesis of both THIQ scaffolds. Namely, inactivation of Adh6 and Ari1 increased production of 4-hydroxyphenethyl-THIQ from ʟ-Hty by 2.0- and 2.4-fold, respectively (**Fig. 4d**). Similarly, inactivation of Adh6 and Ari1 each increased synthesis of phenylpropyl-THIQ from ʟ-APPA each by 1.8-fold (**Fig. 4e**). Collectively, these data demonstrate that that NADPH-dependent oxidoreductases from the Ehrlich pathway limit production of broad THIQ and amphetamine scaffolds from amino acids using our devised retrobiosynthesis approach.

### Extending the substituted amphetamine platform to pharmaceutical products

Previously, it was shown that an *E. coli* strain harboring the ω-transaminase (ω-TA) PP2799 from *Pseudomonas putida* (*Pp*TA) converts exogenously supplemented (*R*/*S*)-PAC to norephedrine^45^ (**Fig. 5a**). Building on this conversion, we sought to streamline norephedrine synthesis by leveraging the ability of our substituted amphetamine platform to efficiently convert ʟ-Phg to PAC *in vivo*. Using literature and homology searches, we selected two additional eukaryotic ω-TA variants to screen, which were sourced from the nematode *Diploscapter pachys* (*Dp*TA) and corn (*Zea mays*; *Zm*TA). ω-TA-encoding genes were chromosomally integrated into a PAC production host possessing three oxidoreductase deletions (*Δadh6 Δari1 Δgre2*) and *Ct*PDC. The resulting strains were grown in 2× SC-URA medium supplemented with benzaldehyde. Under these conditions, HPLC-MS analysis yielded a peak consistent with norephedrine in culture extracts of strains expressing *Pp*TA and *Dp*TA, with *Pp*TA synthesizing 2.1-fold more norephedrine than *Dp*TA (**Supplementary Figure 7**).

**Figure 5.**
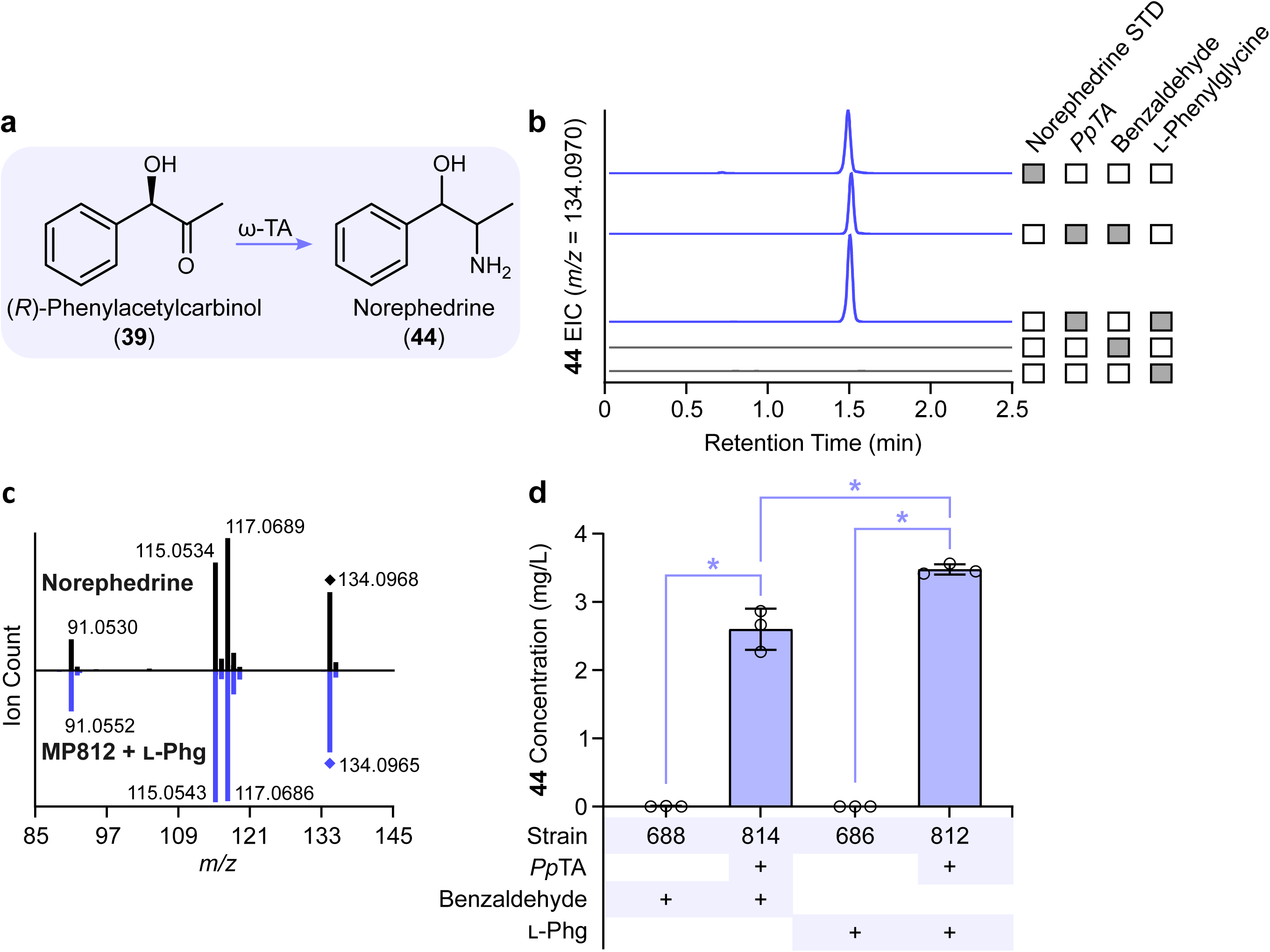
Extending production of a privileged scaffold to an active pharmaceutical ingredient. **a,** Biosynthetic pathway for conversion of (*R*)-phenylacetylcarbinol [(*R*)-PAC] to norephedrine by an ω-transaminase (ω-TA). **b,** Ion-extracted LC-MS chromatogram of norephedrine production strains containing ω-TA from *Pseudomonas putida* (*Pp*TA) supplemented with benzaldehyde (MP814) or ʟ-phenylglycine (MP812). Retention time of yeast-derived norephedrine is compared against an authentic commercial standard. Control chromatograms derived from strains lacking *Pp*TA (MP688 + benzaldehyde; MP686 + ʟ-Phg) are shown for comparison. All strains harbor a pyruvate decarboxylase (PDC) variant from *Candida tropicalis* (*Ct*PDC) and deletions in oxidoreductase-encoding genes (*adh6*Δ *ari1*Δ *gcy1*Δ *gre2*Δ *ypr1*Δ). LC-MS chromatographic experiments were performed three times, with each replicate yielding similar results. **c,** MS/MS fragmentation spectra of commercially-derived norephedrine and spent media from *S. cerevisiae* strain MP812 containing *Pp*TA grown on ʟ-Phg as a sole nitrogen source. MS/MS fragmentation experiments were performed two times, with each replicate yielding similar results. **d,** Comparison of norephedrine titers by engineered yeast strains supplemented with exogenous benzaldehyde or cultivated on ʟ-Phg as a sole source of nitrogen. All strains harbor a pyruvate decarboxylase (PDC) variant from *Candida tropicalis* (*Ct*PDC) and deletions in oxidoreductase-encoding genes (*adh6*Δ *ari1*Δ *gcy1*Δ *gre2*Δ *ypr1*Δ). Asterisk (*) denotes a significant increase (*P* < 0.05) in norephedrine concentration. Error bars represent the mean ± s.d. of *n* = 3 independent biological samples. Statistical differences between control and derivative strains were tested using two-tailed Welch’s *t*-test. Abbreviations: AUC, area under the curve; EIC, extracted ion chromatogram; min, minute; ʟ-Phg, ʟ-phenylglycine; STD, standard; ω-TA, ω-transaminase.

Having confirmed production of norephedrine by *Pp*TA via benzaldehyde supplementation, we next wished to produce norephedrine starting from ʟ-Phg. *Pp*TA was chromosomally integrated into our substituted amphetamine platform possessing five oxidoreductase deletions (*Δadh6 Δari1 Δgcy1 Δgre2 Δypr1*), *Ct*PDC, and *Kd*Aro10. The resulting strain was transformed with pHLUM to restore prototrophy (strain MP812) and grown in 2× YNB medium containing ʟ-Phg as a source of nitrogen. Following HPLC-MS analyses, strain MP812 demonstrated a norephedrine peak (*m/z* – H_2_O = 134.0970 ± 3.7 ppm) that co-eluted with both benzaldehyde-derived norephedrine and a commercial norephedrine standard (**Fig. 5b**). Formation of norephedrine was also dependent on the presence of *Pp*TA. ʟ-Phg-derived norephedrine generated identical fragmentation spectra as an authentic norephedrine standard (**Fig. 5c**), verifying the capacity of our engineered strains to convert ʟ-Phg to norephedrine. Norephedrine synthesis from ʟ-Phg was 34% greater than from supplemented benzaldehyde (**Fig. 5d**), illustrating the tremendous potential of our retrobiosynthetic approach to access structurally and functionally diverse alkaloids.

## DISCUSSION

In this work, we developed a modular and generalizable biocatalytic framework to access structurally and functionally diverse alkaloid scaffolds, including phenyl-THIQs (e.g., solifenacin, cryptostyline), phenethyl-THIQs (e.g., colchicine), and substituted amphetamines (e.g., norephedrine, pseudoephedrine, ephedrine), from simple amino acid precursors. Our novel routes were predicted via retrobiosynthetic logic, a promising yet largely untapped tool in synthetic biology. Retrobiosynthetic analysis begins with target molecule identification—here, privileged alkaloid scaffolds—followed by the prediction of potential precursor molecules and enzymes. Ultimately, a metabolic pathway is assembled in which a series of endogenous or heterologous enzymes catalyze the conversion of a cellular metabolite or chemical building block—in this case, a non-standard amino acid—into the target molecule^46,47^. In our analysis, the Ehrlich pathway was leveraged as a conduit for the bespoke synthesis of target pharmaceutical scaffolds from diverse aldehyde precursors, thereby mirroring the conversion of ʟ-tyrosine into (*S*)-norcoclaurine in traditional yeast-based BIA biosynthesis^16,22^. Other experimentally validated retrobiosynthetic routes include the synthesis of unnatural lactams, 1,4-butanediol, 5-aminolevulinic acid, and short-chain primary amines^48–51^.

Our THIQ-synthesizing strains provide a versatile biocatalytic platform for downstream pharmaceutical production via plant tailoring enzymes or established chemical methods. For example, the (*S*)-phenyl-THIQ core is found in the pharmaceutical solifenacin, which is used to treat overactive bladder and neurogenic detrusor overactivity, and also in cryptostyline natural products^33,52^. Other phenyl-THIQs have promising anti-tumor, anti-HIV, and contraceptive properties^33^. Solifenacin and plant-derived cryptostylines collectively encompass (*S*)-stereoisomers, underscoring the stereochemical advantages of biocatalytic routes to these compounds. In our study, *Tf*NCS-M97V produced enantiopure phenyl-THIQ relative to a racemic standard, yet absolute stereochemistry was not determined due to the absence of an enantiopure (*S*)-phenyl-THIQ standard. Since nearly all NCS variants characterized to date stereospecifically produce (*S*)-THIQs^53^, the enantiopure THIQs produced in our study presumably possess the (*S*)-conformation. Similarly, (*S*)-4-hydroxyphenethyl-THIQ is utilized as a precursor by flowering plants of the Colchicaceae family, such as autumn crocus (*Colchicum autumnale*) and flame lily (*Gloriosa superba*), to synthesize colchicine, which is used in the prevention and treatment of gout^54^. All active NCS variant strains assayed in our study yielded (*S*)-hydroxyphenethyl-THIQ based chiral chromatography of yeast strains harboring *Cj*NCSΔN, which has been shown to stereospecifically produce the (*S*)-conformer^32^. Recently, the complete colchicine biosynthesis pathway was elucidated from *C. autumnale* and is thus primed for reconstruction in our (*S*)-4-hydroxyphenethyl-THIQ host^32,55^. We also extended our retrobiosynthetic framework beyond phenyl- and phenethyl-THIQs to yield phenylpropyl-THIQs starting from ʟ-APPA. Our phenylpropyl-THIQ-producing strain enables pathway extension to THIQs not found in nature, including structures with promising analgesic activities^37^.

To demonstrate the potential of our biocatalytic framework for pharmaceutical manufacturing, we implemented a downstream pathway for production of norephedrine. Production of norephedrine from ʟ-Phg was more efficient than direct supplementation of benzaldehyde used in the industrial synthesis of (*R*)-PAC. In addition to norephedrine, our (*R*)-PAC production strain can be easily adapted to synthesize additional norephedrine isomers, such as norpseudoephedrine, using an (*S*)-selective *Ap*PDC-E469G mutant from *Acetobacter pasteurianus*^56^. Our norephedrine host is also suited for consolidated biocatalytic production of ephedrine and pseudoephedrine through *N*-methylation by newly characterized phenylalkylamine *N*-methyltransferase from *Ephedra sinica* (*Es*PaNMT) or phenylethanolamine *N*-methyltransferase from *Homo sapiens* (*Hs*PNMT)^45,57^. Underscoring the significance of our biocatalytic platform is ephedrine’s inclusion on the World Health Organization’s list of essential medicines, reflecting its role in preventing and treating anesthesia-induced hypotension^58^.

Our bespoke routes to substituted THIQs and amphetamines were initially limited by the availability of Ehrlich pathway-derived aldehyde precursors, which are rapidly converted to fusel alcohols or acids to mitigate their toxic effects^41^. To address this challenge, we blocked the reduction of benzaldehyde to benzyl alcohol via combinatorial deletion of five genes encoding NADPH-dependent aldo-keto reductases (*ADH6*, *ARI1*, *GCY1*, *GRE2*, and *YPR1*), thereby addressing a century-old bottleneck in commercial PAC synthesis^43^. These findings are consistent with previous observations that limited NADPH availability, rather than NADH, decreases benzyl alcohol formation^59^. In addition to the conversion of ʟ-Phg into PAC, we demonstrated that deletion of *ADH6* and *ARI1* broadly improved the conversion of ʟ-Hty and ʟ-APPA into their corresponding (*S*)-4-hydroxyphenethyl- and phenylpropyl-THIQ scaffolds, respectively. The role of these oxidoreductases mirrors that observed in BIA biosynthesis, where many of the same NADPH-dependent enzymes reduce 4-HPAA to tyrosol, thereby impeding production of the committed (*S*)-norcoclaurine scaffold^18^.

Several plant NCS variants were screened to identify optimal candidates to produce each substituted THIQ scaffold. Consistent with prior work, *Tf*NCS-M97V demonstrated exceptional substrate promiscuity, uniquely catalyzing condensation of all aldehydes assayed, and displayed the highest activity with challenging α-substituted aldehydes, namely benzaldehyde from ʟ-Phg and 2-methylbutyraldehyde from ʟ-Val^30,33^. In contrast, *Sd*NCSΔC was the most effective variant for the bulky substrates indole-3-acetaldehyde from ʟ-Trp and 4-phenylbutyraldehyde from ʟ-APPA, while *Cj*NCSΔN exhibited the highest activity on 4-HDCA and 3-phenylpropionaldehyde derived from ʟ-Hty and ʟ-Hph, respectively. *Cj*NCSΔN was recently utilized to reconstitute the heterologous colchicine pathway in *Nicotiana benthamiana* and is the most efficient NCS variant in yeast-based BIA synthesis^18,32,55^. *Cj*NCSΔN has previously been compartmentalized to the peroxisome to improve (*S*)-norcoclaurine synthesis, owing to its toxicity in yeast^60^. Our survey of NCS enzymes also identified *Ac*NCSΔN as a highly active untapped variant with comparable activity to *Cj*NCSΔN on the canonical 4-HPAA substrate. *Ac*NCSΔN exhibited markedly higher specificity for (*S*)-norcoclaurine production relative to *Cj*NCSΔN, illuminating potential application in improving yeast BIA production.

Our work showcases that yeast can derive nitrogen from a broad range of non-standard amino acids, including ʟ-Phg, ʟ-Hty, and ʟ-APPA, thereby establishing an endogenous source of benzaldehyde, 4-HDCA, and 4-phenylbutyraldehyde, respectively. By utilizing the Ehrlich pathway as a biocatalytic entry point, we provide tailored access to these privileged aldehyde precursors without the requisite addition of any heterologous enzymes. As our initial benzaldehyde-producing strain was limited by poor ʟ-Phg utilization and full conversion of ʟ-Phg to (*R*)-PAC is achieved with native yeast enzymes, we screened 36 yeast strains across 11 genera for improved growth on ʟ-Phg and conversion to (*R*)-PAC. Relative to *S. cerevisiae* BY4741, more than 19 yeasts demonstrated superior growth on ʟ-Phg and associated synthesis of PAC and benzyl alcohol. Heterologous expression of the *ARO10* gene encoding phenylpyruvate decarboxylase from *K. dobzhanskii* in our *S. cerevisiae* host improved PAC synthesis 10-fold, underscoring the untapped biocatalytic potential of non-*Saccharomyces* yeasts^39,61^. As our screen focused on strains closely related to *S. cerevisiae*, expanding our search to a broader yeast panel may reveal species with even greater catalytic potential to convert ʟ-Phg to benzaldehyde and PAC. Our findings unveil a new Ehrlich-dependent route to benzaldehyde in engineered yeast and thus provides a foundation for the microbial synthesis of benzaldehyde and benzyl alcohol, which are both key precursors used in the production of food additives, cosmetics, pharmaceuticals, and plastics^62–64^.

Our alkaloid-producing yeast strains are further primed for the *de novo* synthesis of target scaffolds and pharmaceuticals from sugar, as all non-standard amino acids utilized in this study are biosynthesized in nature. ʟ-Phg is produced by *Streptomyces* species as a constituent of the peptide antibiotics virginiamycin S, streptogramin B, and pristinamycin I. ʟ-Phg biosynthesis has also been reconstructed in *E. coli*^65–67^. ʟ-Hty is synthesized by select *Aspergillus* species and cyanobacteria for its use in the production of antifungals (e.g., echinocandin B) and toxins (e.g., microcystins), respectively^68,69^. Similarly, ʟ-Hph is synthesized by cyanobacteria, such as *Nostoc punctiforme* and an *Anabaena* species, whereby the *N. punctiforme* biosynthetic gene cluster has been elucidated and utilized to produce ʟ-Hph from ʟ-Phe in *E. coli*^70,71^. Additionally, *de novo* ʟ-APPA synthesis has been observed by the filamentous fungus *Alternaria alternata*^72^. Incorporation of these non-standard amino acid pathways into engineered strains would enable production of alkaloid scaffolds from sugar, thus precluding the need for amino acid supplementation.

In summary, this work demonstrates a generalizable biocatalytic framework for synthesizing privileged aldehyde-derived pharmaceuticals by rerouting yeast fusel metabolism. We demonstrated the engineering strategy by producing dedicated scaffolds involved in the synthesis of solifenacin, colchicine, and ephedrine—three structurally and functionally diverse alkaloid drugs. We believe that the biocatalytic strategy demonstrated here might be extendable to even more diverse aldehyde intermediates and high-value end-products, including pharmaceuticals and advanced biofuels, from simple amino acid inputs. In a broader context, linking these novel retrobiosynthetic routes to non-standard amino acid biosynthesis and downstream tailoring enzymes or chemical derivatization will enable fully *de novo* production of clinically important pharmaceuticals.

## Acknowledgments

We thank Brad Urquhart and Mark Bernards for assistance with LC-MS analyses and Hannah Lye and Christopher Jia Jun Tran for assistance selecting enzyme candidates. This study was financially supported by the Natural Sciences and Engineering Research Council of Canada (NSERC) through a CGS-M scholarship to A.E.C.R. and research grants to M.E.P. (RGPIN-2023-04542 and DGECR-2023-00416). M.A.L. and C.A.R. acknowledge financial support from the Natural Sciences and Engineering Research Council of Canada (NSERC) and the Conselho Nacional de Desenvolvimento Científico e Tecnológico, Brazil (CNPq), respectively.

## Author contributions

M.E.P., M.A.L., and A.E.C.R. designed the research. A.E.C.R., D.K., J.R.G., M.H., and L.B. performed the experiments with supervision from M.E.P. and M.A.L. C.A.R. and M.A.L. selected and provided non-*Saccharomyces* yeast species for screening. A.E.C.R. and M.E.P. wrote the manuscript. All authors edited the manuscript.

## Additional information

Supplementary information is available in the online version of the paper. Correspondence and requests for materials should be addressed to M.E.P.

## Data availability

Data supporting the findings of this work are available within the paper and its SI Appendix. Source data for the figures in this paper and the SI Appendix are provided in Dataset S1.

## METHODS

### Chemicals and reagents

General chemicals and reagents were purchased commercially, including ʟ-2-amino-5-phenyl-pentanoic acid (AmBeed), benzaldehyde (MilliporeSigma), benzyl alcohol (MilliporeSigma), dopamine hydrochloride (MilliporeSigma), ʟ-homophenylalanine (Thermo Fisher Scientific), ʟ-homotyrosine hydrobromide (Chem-Impex), 4-hydroxydihydrocinnamaldehyde (4-HDCA, AmBeed), isovaleraldehyde (MilliporeSigma), methional (MilliporeSigma), (*R,S*)-norcoclaurine hydrochloride (MilliporeSigma), norephedrine hydrochloride (MilliporeSigma), 2-phenylacetaldehyde (MilliporeSigma), (*R*)-phenylacetylcarbinol (Toronto Research Chemicals), 4-phenylbutyraldehyde (Toronto Research Chemicals), ʟ-2-phenylglycine (MilliporeSigma), 3-phenylpropionaldehyde (Thermo Fisher Scientific), sodium pyruvate (BioBasic). The yeast MoClo toolkit (YTK) was purchased from Addgene (kit #1000000061)^73^.

### Plasmids, strains, and growth media

The quadruple auxotrophic *S. cerevisiae* strain BY4741 (*MATa his3Δ1 leu2Δ0 met15Δ0 ura3Δ0*) is the origin of all engineered strains in this study. Yeast cultures were routinely grown in YPD (10 g/L Bacto Yeast Extract, 20 g/L Bacto peptone, 20 g/L glucose), Yeast Nitrogen Base (YNB) medium (6.7 g/L Difco YNB without amino acids and 20 g/L glucose or sucrose), or synthetic complete (SC) medium (6.7 g/L Difco YNB without amino acids, 1.62-1.92 g/L Drop-out Medium Supplements without appropriate amino acids, and 20 g/L glucose or sucrose). Selection using auxotrophic markers was performed in SC medium lacking uracil or histidine, or in YNB medium without amino acids. Non-*Saccharomyces* yeasts were maintained on YM or YPD medium. Strains grown on individual amino acids as a sole nitrogen source were cultivated in 2× YNB without ammonium sulfate and without amino acids (1.7 g/L) containing 20 g/L glucose. Oligonucleotides utilized in this study are listed in **Supplementary Data 1**. Plasmids employed in this work are provided in **Supplementary Table 4** and engineered *S. cerevisiae* strains are provided in **Supplementary Data 2**.

### Yeast strain construction

Genetic modifications to yeast were made using Golden Gate cloning and the Yeast MoClo Toolkit or via CRISPR-Cas9 and *in vivo* DNA assembly. *NCS* (911b locus), *PDC* (106a locus), and *ARO10* (308a locus) gene variants were chromosomally integrated using CRISPR-Cas9. Cas9 and guide RNA (gRNA) were delivered to yeast using pBBK94 and pBBK95^74^. gRNAs are provided in **Supplementary Table 5** and were assembled by oligo annealing and extension and assembled *in vivo* using gap repair with BsaI- and NotI-digested pBBK94 and pBBK95^74^. *S. cerevisiae* was transformed using 50 μL PEG, lithium acetate, single-stranded salmon sperm DNA reactions via heat shock at 42 °C for 30 minutes. Cells were recovered in YPD (antibiotic selection) or SC medium (auxotrophic selection) for 16-20 h without shaking at 30 °C. Chromosomal loci for integration were selected based on a prior study^75^. Synthetic DNAs employed in this study are provided in **Supplementary Data 3**.

Oxidoreductase gene deletions were performed using CRISPR-Cas9 via plasmids pBBK94 and pBBK95^74^. Genes targeted for deletion were replaced by a synthetic 23 bp Cas9 target site (T9) for subsequent DNA integrations. Design of all gRNAs was performed by cross-referencing both CCTop^76^ and CRISPRi^77^ tools. Cassettes and pathways for chromosomal integration of a dopamine biosynthesis pathway (*PpDODC*, *BvCYP76AD5*, *ScARO4^K229L^*, *ScARO7^G141S^*, *ScTYR1*, and *ScURA3*) and ω-transaminases (*PpTA*, *DpTA*, and *ZmTA*) were assembled using the yeast MoClo toolkit^73^.

### Growth curves

Growth curves were generated in triplicate in 96-well microtiter plates containing 180 µl of medium. To assess utilization of amino acids as sources of nitrogen, prototrophic engineered strains and non-conventional yeasts were cultivated in 2× YNB medium without ammonium sulfate and without amino acids supplemented with 0.5-1.0 g/L of the target amino acid and 2% glucose. All cultures were inoculated using saturated overnight cultures to an initial OD_600_ of 0.1-0.3 and sealed with a Breathe-Easy sealing film (Diversified Biotech). Absorbance was measured at 600 nm every 20 minutes with a BioTek Epoch 2 microplate spectrophotometer (Agilent Technologies) over the course of 120 hours. Maximum specific growth rates (*u*_max_) and area under the curve (AUC) were determined from triplicate cultures using BioTek Gen6 data analysis software (Agilent Technologies).

### Microtiter plate assay for production of heterologous metabolites

Non-conventional yeast and engineered *S. cerevisiae* strains were picked in triplicate into deep 96-well plates containing 0.5 ml of 2× SC medium containing 2% glucose or sucrose, or 2× YNB medium without amino acids and containing 5 g/l ammonium sulfate and 2% glucose or sucrose. Cultures were grown overnight, diluted 50× into fresh YNB medium without ammonium sulfate and without amino acids supplemented with 2% glucose and 0.5-1.0 g/l of a target amino acid as nitrogen source. For benzaldehyde supplementation, overnight cultures were diluted 50× into fresh 2× SC medium containing 2% glucose, 7 mM benzaldehyde, and 8.4 mM pyruvate. Benzaldehyde and pyruvate were supplemented every 24 hours for a total of 96 hours using a 10× feed solution containing 2× SC medium, 70 mM benzaldehyde, and 84 mM pyruvate. All deep-well plates were incubated at room temperature with shaking for 72-96 hours and samples were frozen at - 20 °C prior to LC-MS analysis.

### Chemical synthesis of substituted tetrahydroisoquinolines

Chemical standards of substituted tetrahydrosioquinoline standards were prepared via phosphate-catalyzed reaction of dopamine with various aldehydes according to a previous report^33^. Ten ml reactions contained 10 mM dopamine, 20 mM aldehyde, and 10 mM sodium ascorbate in 300 mM pH 6 potassium phosphate buffer with 50% acetonitrile. Reactions were incubated at 60 °C for 16-18 hours. Serial dilutions of THIQ stock standards were prepared in 30% acetonitrile and frozen at −20 °C prior to LC-MS analysis.

### LC-MS, HPLC-UV, and chiral analysis of heterologous metabolites

LC-MS was utilized to detect substituted THIQs, (*S*)-norcoclaurine, (*R*)-PAC, and norephedrine from yeast cultures grown in 96-well plates. To extract metabolites, 25 µl cell broth was combined with 60 µl 100% acetonitrile in a 96-well V-bottom microtiter plate and samples were shaken vigorously for five minutes. Subsequently, 115 µl of sterile water was added to give a final volume of 200 µl in 30% and an eight-fold dilution of cultures. Extracts were centrifuged for 20 minutes at 4,000 RCF and supernatants were transferred to glass vials with inserts prior to LC-MS analysis.

Unless stated otherwise, LC-MS samples were analyzed using a Waters ACQUITY UPLC I-Class system coupled to a Waters Xevo^TM^ G2-S QToF mass spectrometer. Five µl of extract or standard was injected and separated using a Waters ACQUITY UPLC HSS T3 column (100 mm × 2.1 mm; 1.8 µm). Metabolites were separated using the following gradient: 10-85% B from 0 to 6 minutes, held at 85% B from 6 to 7 minutes, and returned to 10% B from 7 to 7.1 minutes. The flow rate was maintained at 0.3 ml/minute. Solvent A was 0.1% formic acid and solvent B was 0.1% formic acid in 100% acetonitrile. The source conditions were set as follows: capillary voltage, 2.0 kV; cone voltage, 40 V; source temperature, 150 °C; desolvation gas flow, 1,000 l/h; and cone gas flow, 50 l/h. Lockmass correction was applied using a continuous infusion of leucine-enkephalin (500 ng/ml) as reference. The lockspray reference ion (*m/z* = 556.2771 in positive mode) was monitored every ten seconds with a scan time of 0.3 seconds, averaged over three scans, to ensure mass accuracy throughout the run. MS/MS analysis was performed using a collision energy ramp of 15-50 V.

In other experiments, LC-MS analysis was performed using an Agilent 1260 LC system (Agilent Technologies) equipped with an Eclipse Plus RRHT C-18 column (3.0 × 100 mm, 1.8 µm; Agilent Technologies). Twenty µl of extract or standard was injected and separated using the following gradient: 15% B from 0 to 7 minutes, 15-90% B from 7 to 11 minutes, and return to 15% B from 11 to 13 minutes, followed by a 10-minute post-run equilibration. The flow rate was maintained at 0.25 ml/minute. Solvent A was 0.1% formic acid and solvent B was 0.1% formic acid in 90% acetonitrile. ESI-TOF parameters were as follows: drying gas at 350 °C, 10 mL/min; nebulizer at 45 PSI; VCap at 4000 V; fragmentor at 130 V. Spectra were collected at 1.03/s (9729 transients/spectrum) in the 100–1700 *m*/*z* range. Reference mass solution (121.050873 *m*/*z* and 922.009798 *m*/*z*) was infused constantly via a second nebulizer at 15 psi.

Benzyl alcohol was detected using HPLC-UV at 263 nm and the above Agilent 1260 LC system (Agilent Technologies) with identical sample preparation, column, and separation conditions.

Chiral separation of (*R,S*)-norcoclaurine and chemically-synthesized (*R,S*)-THIQs was performed as described previously^18,78^. Culture supernatants, commercial (*R,S*)-norcoclaurine, or chemically-synthesized (*R,S*)-THIQs were diluted in water and 10 µl was loaded onto a Shodex ORPak CDBS-453 chiral column (4.6 × 150 mm). Chiral compounds were separated using an Agilent 1260 LC system (Agilent Technologies) and the following gradient: 0-13 min, 5% B; 14-17 min, 95% B; and 18-30 min, 5% B, followed by a 10-minute post-run equilibration, where solvent A was 0.1% formic acid and solvent B was 0.1% formic acid in 90% acetonitrile. Flow rate was 0.25 ml/min and temperature was held at 25 °C. A modified gradient was employed for chiral resolution of (*R,S*)-phenylpropyl-THIQ (**31**) using the same solvents, temperature, and flow rate: 0-23 min, 5% B; 24-27 min, 95% B; and 28-30 min, 5% B, followed by a 10 minute post-run equilibration. Retention time of yeast-derived norcoclaurine was compared to a commercial (*R,S*)-norcoclaurine standard. Retention time of other substituted THIQs were compared to racemic substituted THIQ standards prepared through chemical condensation of dopamine and corresponding aldehydes.

### Statistical analyses

All numerical values are depicted as means ± s.d. Statistical differences between control and engineered strains were assessed via two-tailed Welch’s *t*-tests assuming equal variances. In all cases, *P*-values < 0.05 were considered significant.

