## Supplementary Information for "An Ehrlich-inspired retrobiosynthesis of pharmaceutical scaffolds"

**SUPPLEMENTARY FIGURES**

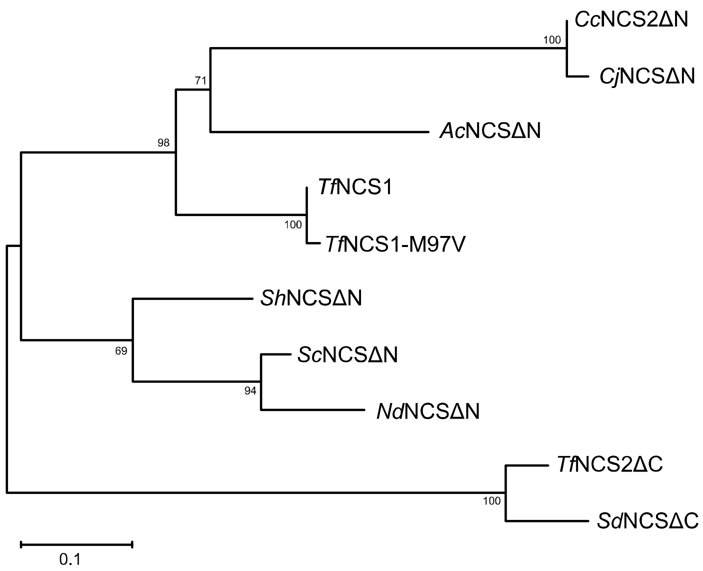


**Supplementary Figure 1. Phylogenetic analysis of norcoclaurine synthase (NCS) variants employed in this study.** NCS protein sequences were aligned using MUSCLE and the maximum-likelihood tree was constructed using MEGA11[1].

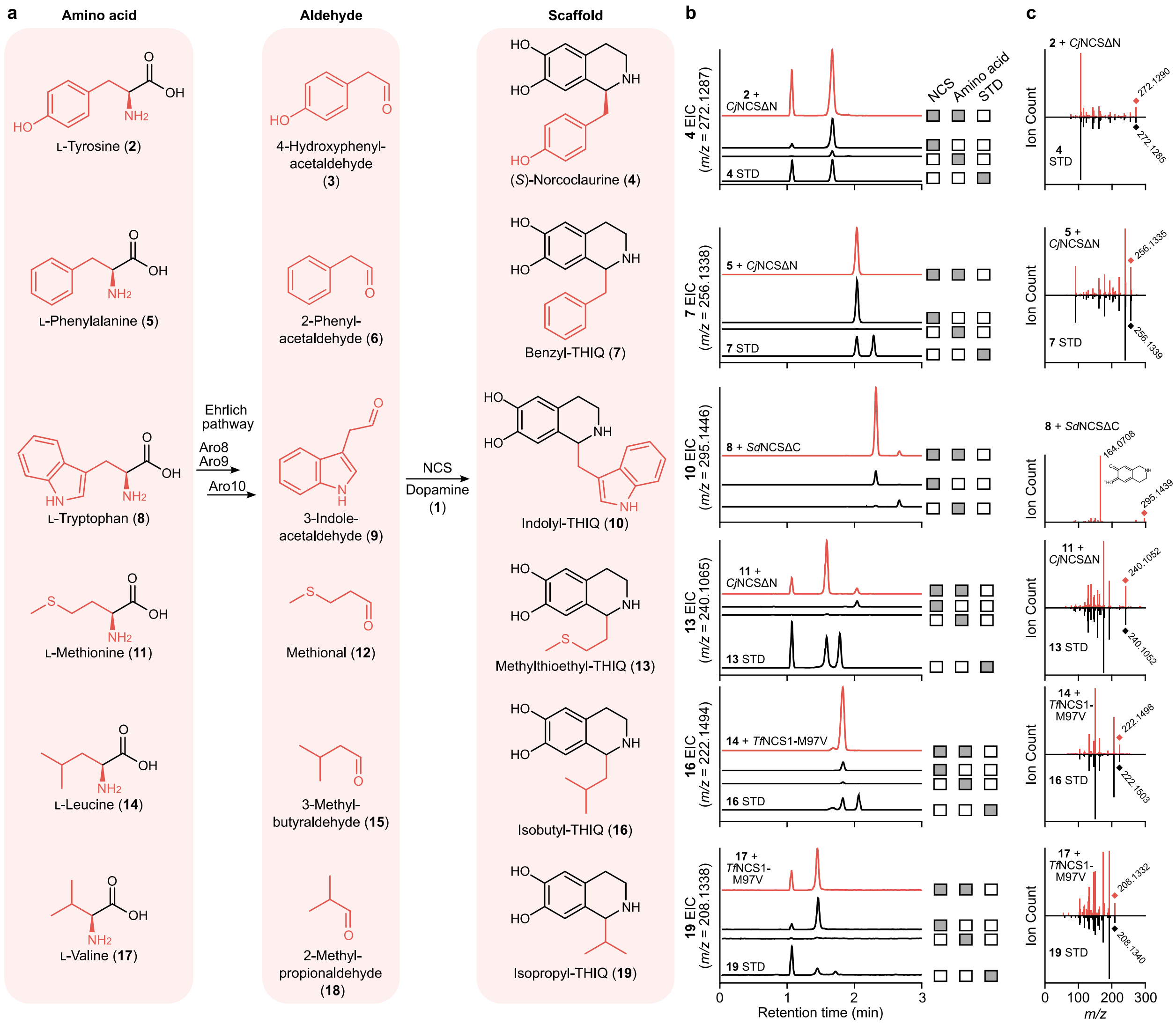


**Supplementary Figure 2. Characterization of substituted tetrahydroisoquinolines (THIQs) derived from endogenous amino acids.** Biosynthetic pathways (**a**), ion-extracted LC-MS chromatograms (**b**), and MS/MS fragmentation spectra (**c**) of substituted THIQs derived from the yeast Ehrlich pathway. Engineered yeast strains were grown on a mixture of Ehrlich pathway amino acids [L-Tyr (**2**), L-Phe (**5**), L-Trp (**8**), L-Met (**11**), L-Leu (**14**), and L-Val (**17**)] as a source of nitrogen to promote formation of substituted THIQs. LC-MS chromatogram experiments included controls lacking an NCS biosynthetic enzyme or with ammonium sulfate in place of exogenous Ehrlich amino acids. Yeast-derived THIQs were compared against chemical standards attained from a commercial supplier [(*R*,*S*)-norcoclaurine (**4**)] or derived through the phosphate-catalyzed condensation (**7**, **13**, **16**, **19**) of dopamine (**1**) and corresponding aldehydes (**6**, **12**, **15**, **18**). A commercial source of indolyl-THIQ (**10**) and its corresponding aldehyde (**9**) was not available, and fragmentation of **10** was predicted using CFM-ID[2]. Incorporation of amino acids into aldehydes and substituted THIQs is shown in red. LC-MS chromatographic and MS/MS fragmentation experiments were performed three times and two times, respectively, with each replicate yielding similar results. Abbreviations: EIC, extracted ion chromatogram; min, minute; NCS, norcoclaurine synthase; STD, standard; THIQ, tetrahydroisoquinoline.

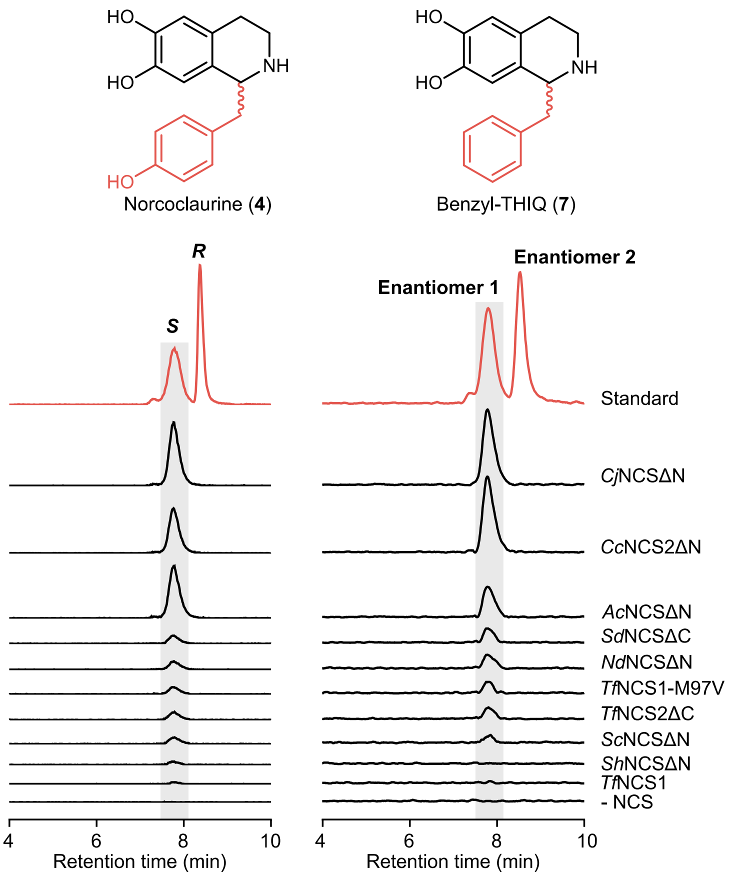


**Supplementary Figure 3. Chiral analysis of substituted tetrahydroisoquinolines (THIQs) produced by engineered NCS yeast strains.** Yeast-derived norcoclaurine (**4**) was compared against commercial (*R*,*S*)-norcoclaurine standard. *Cj*NCS has previously been shown to stereospecifically produce (*S*)-norcoclaurine[3]. Yeast-derived benzyl-THIQ (**7**) was compared against authentic (*R*,*S*)-(**7**) standard prepared by the phosphate-catalyzed condensation of dopamine (**1**) and 2-phenylacetaldehyde (**6**). Enantiopure yeast-derived conformers are shaded. Absolute stereochemistry of yeast-derived benzyl-THIQ (**7**) could not be assigned due to the absence of an authentic enantiopure standard. Chiral LC-MS chromatographic experiments were performed three times with each replicate yielding similar results. Abbreviations: min, minute; NCS, norcoclaurine synthase; THIQ, tetrahydroisoquinoline.

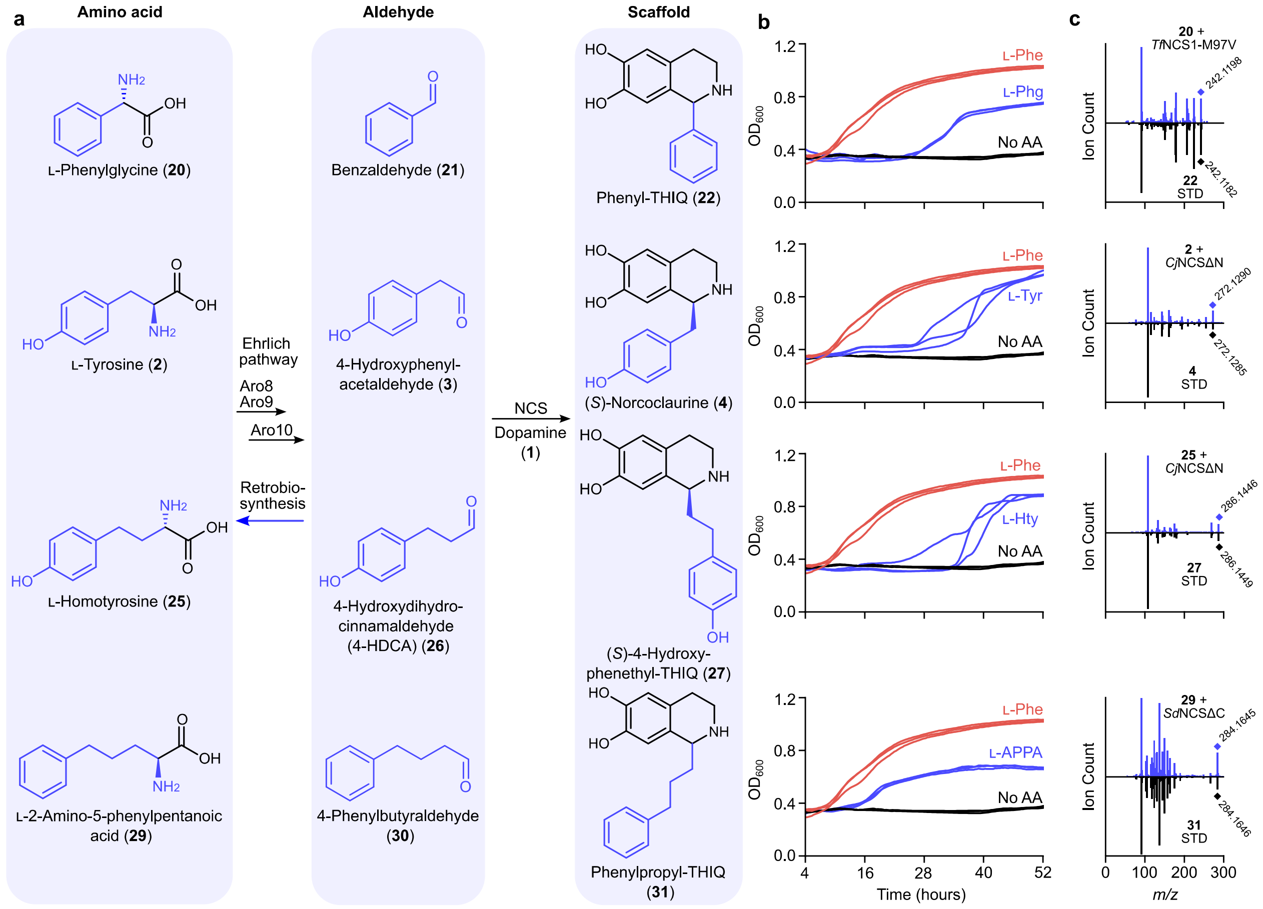


**Supplementary Figure 4. Characterization of pharmaceutical tetrahydroisoquinolines (THIQs) derived from non-standard amino acids.** Proposed formation (**a**), non-standard amino acid utilization (**b**), and MS/MS fragmentation spectra (**b**) involved in production of substituted THIQs. Engineered yeast strains were grown on non-standard amino acids [L-Phg (**20**), L-Hty (**25**), or L-2-amino-5-phenylpentanoic acid (L-APPA, **29**)] as a sole source of nitrogen. For growth curves, growth on non-standard amino acids was compared against L-Phe (**5**) (red) and control cultures lacking nitrogen (no AA; black) where $n = 3$ independent biological replicates are overlaid for each condition. Fragmentation of yeast-derived THIQs (**22**, **27**, **31**) were compared against chemical standards derived from the phosphate-catalyzed condensation of dopamine (**1**) and corresponding aldehydes (**21**, **26**, **30**). Incorporation of amino acids into aldehydes and substituted THIQs is shown in blue. MS/MS fragmentation experiments were performed two times, yielding similar results. Abbreviations: AA, amino acid; L-APPA, L-2-amino-5-phenylpentanoic acid; L-Hty, L-homotyrosine; NCS, norcoclaurine synthase; L-Phe, L-phenylalanine; L-Phg, L-phenylglycine; STD, standard; THIQ, tetrahydroisoquinoline; L-Tyr, L-tyrosine.

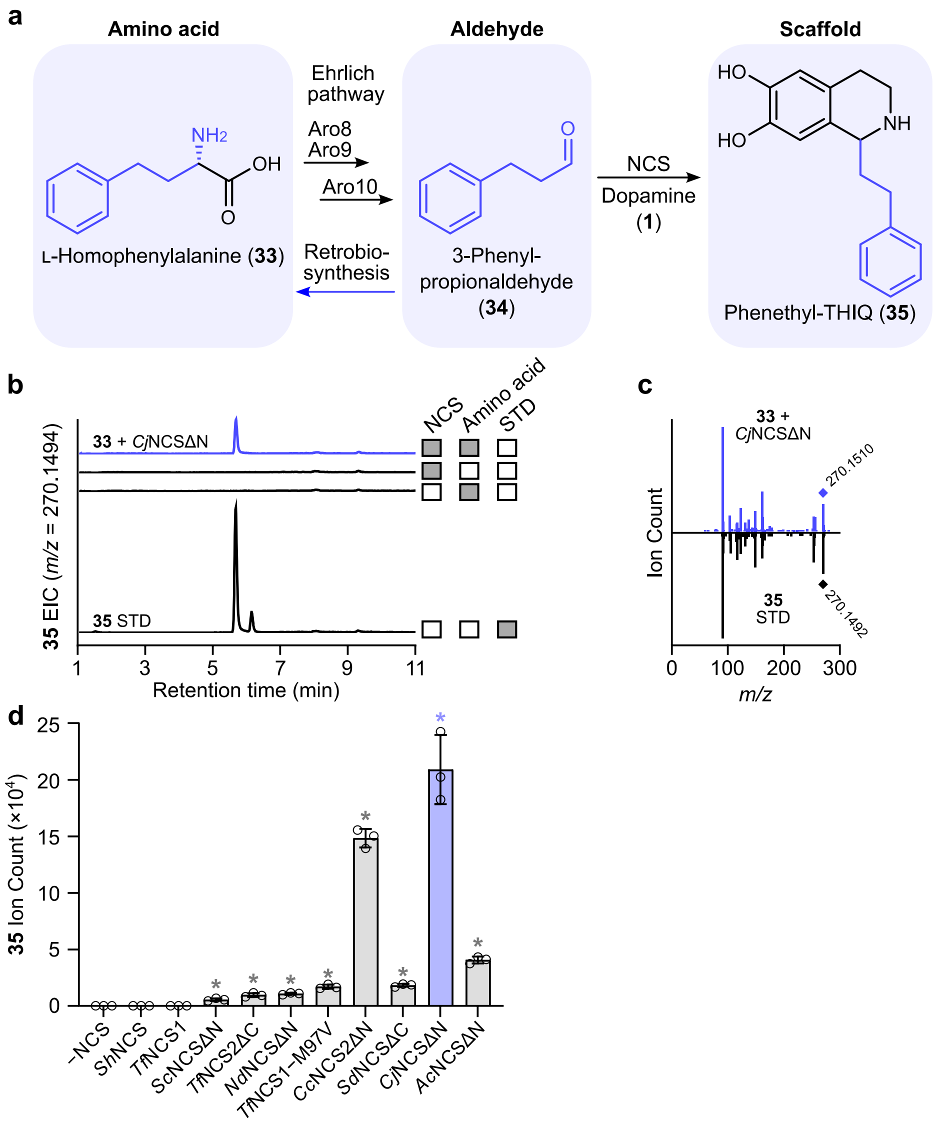


**Supplementary Figure 5. Characterization of phenethylisoquinoline (phenethyl-THIQ) derived from L-homophenylalanine (L-Hph).** Proposed formation (**a**), ion-extracted LC-MS chromatogram (**b**), MS/MS fragmentation spectra (**c**), and quantitative production (**d**) of phenethyl-THIQ derived from the yeast Ehrlich pathway. Engineered yeast strains were grown on L-Hph (**33**) as a source of nitrogen to promote formation of phenethyl-THIQ (**35**). Controls lacking an NCS biosynthetic enzyme or with ammonium sulfate in place of L-Hph were included. Yeast-derived phenethyl-THIQ was compared against a chemical standard derived from the phosphate-catalyzed condensation of dopamine (**1**) and 3-phenylpropanal (**34**). Incorporation of L-Hph into 3-phenylpropanal and phenethyl-THIQ is shown in blue. LC-MS analysis was performed on an Agilent 1260 LC system equipped with a C-18 Eclipse Plus RRHT column (3.0 × 100 mm, 1.8 µm; Agilent Technologies). LC-MS chromatographic and MS/MS fragmentation experiments were performed three times and two times, respectively, with each replicate yielding similar results. For comparison of engineered NCS strains, error bars represent the mean ± s.d. of $n = 3$ independent biological replicates. Abbreviations: EIC, extracted ion chromatogram; L-Hph, L-homophenylalanine; min, minute; NCS, norcoclaurine synthase; STD, standard; THIQ, tetrahydroisoquinoline.

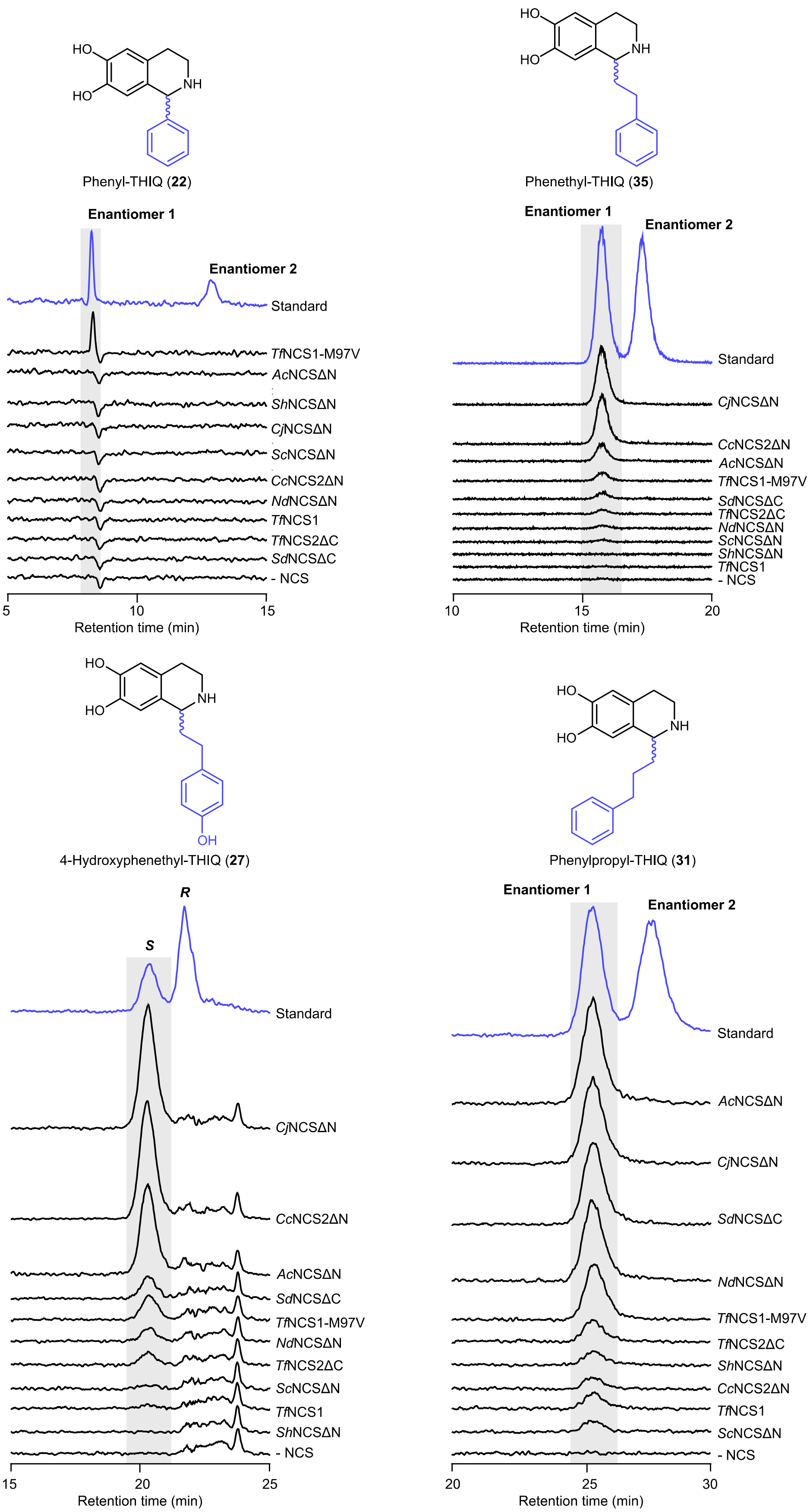

HO
HO
NH
Phenyl-THIQ (22)
Enantiomer 1
Enantiomer 2
Standard
TfNCS1-M97V
AcNCSΔN
ShNCSΔN
CjNCSΔN
ScNCSΔN
CcNCS2ΔN
NdNCSΔN
TfNCS1
TfNCS2ΔC
SdNCSΔC
- NCS
5
10
15
Retention time (min)
Phenethyl-THIQ (35)
Enantiomer 1
Enantiomer 2
Standard
CjNCSΔN
CcNCS2ΔN
AcNCSΔN
TfNCS1-M97V
SdNCSΔC
TfNCS2ΔC
NdNCSΔN
ScNCSΔN
ShNCSΔN
TfNCS1
- NCS
10
15
20
Retention time (min)
OH
4-Hydroxyphenethyl-THIQ (27)
S
R
Standard
CjNCSΔN
CcNCS2ΔN
AcNCSΔN
SdNCSΔC
TfNCS1-M97V
NdNCSΔN
TfNCS2ΔC
ScNCSΔN
TfNCS1
ShNCSΔN
- NCS
15
20
25
Retention time (min)
Phenylpropyl-THIQ (31)
Enantiomer 1
Enantiomer 2
Standard
AcNCSΔN
CjNCSΔN
SdNCSΔC
NdNCSΔN
TfNCS1-M97V
TfNCS2ΔC
ShNCSΔN
CcNCS2ΔN
TfNCS1
ScNCSΔN
- NCS
20
25
30
Retention time (min)

**Supplementary Figure 6. Chiral analysis of substituted tetrahydroisoquinolines (THIQs) derived from non-standard amino acids.** Yeast-derived THIQs (**22, 27, 31, 35**) were compared against (*R,S*)-THIQ standards prepared from the phosphate-catalyzed condensation of dopamine (**1**) and corresponding aldehydes (**21, 26, 30, 34**). Enantiopure yeast-derived conformers are shaded. *Cj*NCS has previously been shown to stereospecifically produce (*S*)-4-hydroxyphenethyl-THIQ (**27**)[4]. Absolute stereochemistry of other yeast-derived THIQs (**22, 31, 35**) could not be assigned due to the absence of enantiopure standards. Chiral LC-MS chromatographic experiments were performed three times with each replicate yielding similar results. Abbreviations: min, minute; NCS, norcoclaurine synthase; THIQ, tetrahydroisoquinoline.

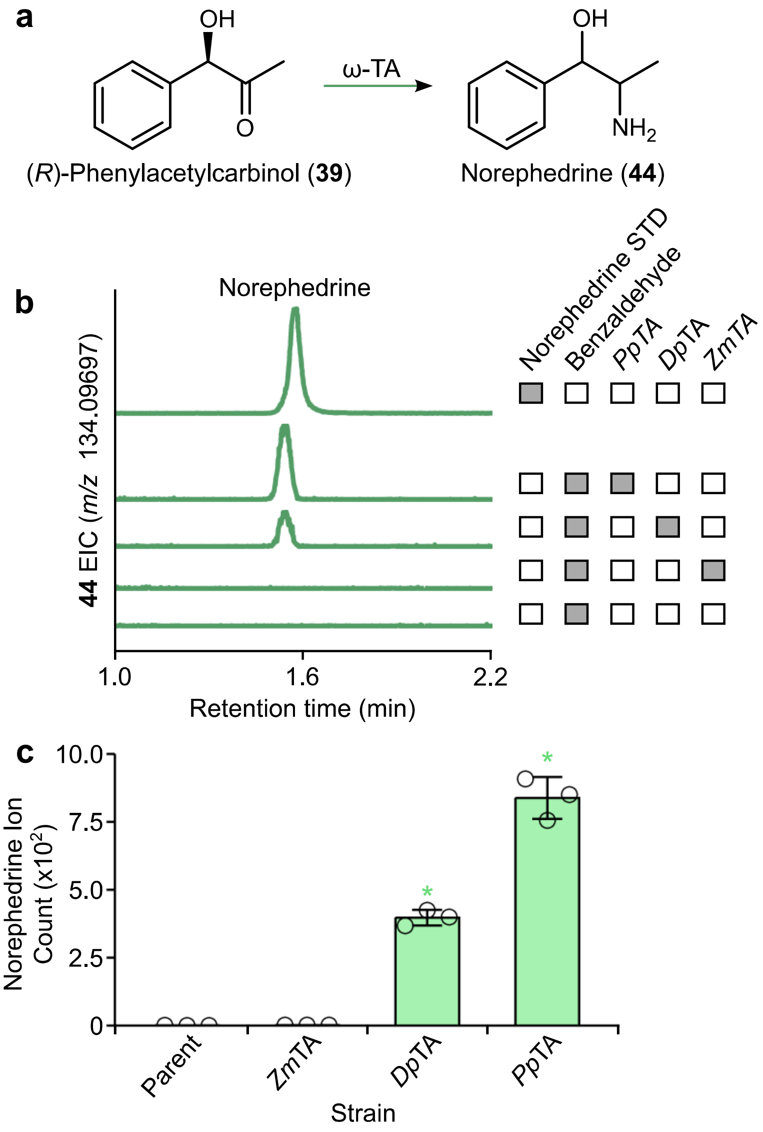


**Supplementary Figure 7. Screening ω-transaminases (ω-TAs) for conversion of (*R*)-phenylacetylcarbinol to norephedrine. a,** Biosynthetic pathway for conversion of (*R*)-phenylacetylcarbinol [(*R*)-PAC] to norephedrine by an ω-transaminase (ω-TA). **b,** Ion-extracted LC-MS chromatogram of strain MP510 (*Ct*PDC, *adh6*Δ *ari1*Δ *gre2*Δ) harboring different ω-TA variants supplemented with benzaldehyde. Retention time of yeast-derived norephedrine is compared against an authentic commercial standard. A control chromatogram derived from strain MP510 lacking an ω-TA enzyme is shown for comparison. LC-MS chromatographic experiments were performed three times, with each replicate yielding similar results. **c,** Norephedrine production by different ω-TA enzyme variants. Production of norephedrine from exogenous benzaldehyde by three ω-TA variants was compared against a parent control (MP510) lacking a ω-TA variant. Asterisk (*) denotes a significant increase ($P < 0.05$) in norephedrine concentration relative to the MP510 parent strain. Error bars represent the mean ± s.d. of $n = 3$ independent biological samples. Statistical differences between control and derivative strains were tested using two-tailed Welch's *t*-test. Abbreviations: EIC, extracted ion chromatogram; min, minute; STD, standard; ω-TA, ω-transaminase.

### SUPPLEMENTARY TABLES

**Supplementary Table 1. Norcoclaurine synthase (NCS) candidates characterized or utilized in this study.**

| Enzyme | Truncation | Species | % Identity[a] | Accession | Reference |
|---|---|---|---|---|---|
| *Ac*NCSΔN | $\Delta N_{25}$ | *Aquilegia coerulea* | 63.5 | PIA58613.1 | This study |
| *Cc*NCS2ΔN | $\Delta N_{34}$ | *Coptis chinenis* | 97.5 | KAF9595526.1 | This study |
| *Cj*NCSΔN | $\Delta N_{34}$ | *Coptis japonica* | 100 | A2A1A1.2 | [3] |
| *Nd*NCSΔN | $\Delta N_{19}$ | *Nandina domestica* | 56.7 | NA | [5] |
| *Sc*NCSΔN | $\Delta N_{19}$ | *Sanguinaria canadensis* | 54.5 | NA | [5] |
| *Sd*NCSΔC | $\Delta C_{25}$ | *Stylophorum diphyllum* | 44.9 | NA | [5] |
| *Sh*NCSΔN | $\Delta N_{19}$ | *Sinopodophyllum hexandrum* | 60.1 | AIT42265.1 | [6] |
| *Tf*NCS1 | - | *Thalictrum flavum* | 63.5 | ACO90247.1 | [7] |
| *Tf*NCS1-M97V | $\Delta N_{7}$ | *Thalictrum flavum* | 63.5 | 6Z82_A | [8] |
| *Tf*NCS2ΔC | $\Delta C_{25}$ | *Thalictrum flavum* | 48.1 | NA | [5] |

[a] Relative to *Cj*NCSΔN

**Supplementary Table 2. Non-conventional yeast species employed in this study.**

| Species | Strain ID |
|---|---|
| *Kazachstania bromeliacearum* | UFMG-CM-Y35 |
| *Ka. humilis* | CBS 5658 |
| *Kluyveromyces aestuarii* | CBS 4904 |
| *Kl. bulgaricus* | CBS 5829 |
| *Kl. dobzhanskii* | CBS 5062 |
| *Kluyveromyces* hybrid[a] | CBS 6925 |
| *Kl. lactis* | CBS 4693 |
| *Kl. marxianus* | CBS 6923 |
| *Kl. nonfermentans* | CBS 8778 |
| *Kl. starmeri* | UFMG-CM-Y-3682 |
| *Kl. starmeri* | UFMG-CM-Y7764 |
| *Lachancea kluyveri* | NRRL Y 12651 |
| *Lachancea* sp. *"fantastica"* | CBS 6924 |
| *L. nothofagi* | UFMG-YM1.2 |
| *L. nothofagi* | UFMG-YM2.2 |
| *L. thermotolerans* | CBS 6340 |
| *Nakaseomyces glabratus* | CBS 138 |
| *N. glabratus* | UCDFST Y55 |
| *Pichia* sp. | UFMG-CM-314 |
| *Pichia* sp. | UFMG-CM-Y7765 |
| *Saccharomyces cerevisiae* | A2.9 |
| *S. paradoxus* | CBS 432 |
| *Tetrapisispora blattae* | CBS 6285 |
| *Torulaspora delbrueckii* | 1996 89A |
| *To. delbrueckii* | 1996 109A |
| *To. delbrueckii* | 1996 143A |
| *To. delbrueckii* | 1996 169 |
| *To. delbrueckii* | UFMG-L-630 |
| *Vanderwaltozyma yarrowii* | CBS 2684 |
| *V. yarrowii* | CBS 6070 |
| *V. yarrowii* | CBS 8242 |
| *Zygosaccharomyces machadoi* | UFMG-CM-Y47 |
| *Zs. rouxii* | UFMG-CM-Y7766 |
| *Zs. rouxii* | UFMG-CM-Y6367 |
| *Zygotorulaspora cariocana* | UFMG-CM-Y6340 |
| *Zt. cariocana* | UFMG-CM-Y6404 |

[a] *Kluyveromyces marxianus* and *Kluyveromyces thermotolerans*[9]

**Supplementary Table 3. Conversion of L-phenylglycine to benzyl alcohol and phenylacetylcarbinol by non-conventional yeast species**

| Species | Strain ID | Phenylacetylcarbinol abundance[a] | Benzyl alcohol abundance[b] |
|---|---|---|---|
| *Kluyveromyces aestuarii* | CBS 4904 | 0 | 65.6 ± 0.4 |
| *Kl. bulgaricus* | CBS 5829 | 60802 ± 3179 | 6.0 ± 0.8 |
| *Kl. dobzhanskii* | CBS 5062 | 257060 ± 14132 | 62.3 ± 5.5 |
| *Kl. marxianus* | CBS 6923 | 0 | 52.8 ± 4.2 |
| *Kl. nonfermentans* | CBS 8778 | 0 | 20.9 ± 0.9 |
| *Kl. starmeri* | UFMG-CM-Y7764 | 0 | 18.4 ± 1.3 |
| *Lachancea kluyveri* | NRRL-Y-12651 | 0 | 30.6 ± 6.4 |
| *Lachancea* sp. *"fantastica"* | CBS 6924 | 0 | 4.7 ± 0.4 |
| *L. nothofagi* | UFMG-YM1.2 | 0 | 10.7 ± 0.2 |
| *L. nothofagi* | UFMG-YM2.2 | 0 | 11.1 ± 1.1 |
| *L. thermotolerans* | CBS 6340 | 0 | 6.5 ± 0.6 |
| *Nakaseomyces glabratus* | CBS 138 | 6204 ± 200 | 42.1 ± 1.4 |
| *N. glabratus* | UCDFST Y55 | 0 | 29.3 ± 1.5 |
| *Pichia* sp. | UFMG-CM-Y7765 | 0 | 1 ± 0.2 |
| *Pichia* sp. | UFMG-CM-Y314 | 0 | 2.5 ± 0.7 |
| *Saccharomyces cerevisiae* | BY4741 | 19842 ± 3317 | 6.2 ± 0.1 |
| *S. cerevisiae* | A2.9 | 43393 ± 4860 | 5 ± 0.5 |
| *S. paradoxus* | CBS 432 | 101130 ± 1438 | 8 ± 0.1 |
| *Torulaspora delbrueckii* | 1996 89A | 0 | 5.6 ± 0.1 |
| *To. delbrueckii* | 1996 109A | 0 | 2.4 ± 0.5 |
| *To. delbrueckii* | 1996 143A | 0 | 4.7 ± 0.1 |
| *To. delbrueckii* | 1996 169 | 0 | 4.6 ± 0.6 |
| *To. delbrueckii* | UFMG-L-630 | 0 | 9 ± 0.5 |
| *Vanderwaltozyma yarrowii* | CBS 2684 | 82400 ± 4522 | 5.4 ± 0.6 |
| *V. yarrowii* | CBS 6070 | 0 | 25.6 ± 6.3 |
| *Zygosaccharomyces machadoi* | UFMG-CM-Y47 | 0 | 1.3 ± 0.4 |
| *Zs. rouxii* | UFMG-CM-Y7766 | 0 | 5.3 ± 0.3 |
| *Zygotorulaspora cariocana* | UFMG-CM-Y6404 | 77495 ± 1222 | 11.9 ± 0.4 |
| *Zt. cariocana* | UFMG-CM-Y6340 | 48314 ± 4328 | 13.3 ± 1.0 |

[a] Detected with LC-MS
[b] Detected with HPLC-UV

**Supplementary Table 4. Plasmids employed in this study.**

| **Plasmid** | **Description** | **Source or reference** |
|---|---|---|
| pBBK94 | pCAS-Tyr-[gRNA: 2x BsaI+NotI cut sites] ($Kan^R$; $G418^R$) | [10] |
| pBBK95 | pCAS-Tyr-[gRNA: 2x BsaI+NotI cut sites] ($Hyg^R$) | [11] |
| pHLUM | *HIS3, LEU2, URA3, MET15* complementation plasmid | [12] |
| pYTK-PP-*ScARO4$^{FBR}$* | *ScARO4$^{FBR}$* part plasmid | This study |
| pYTK-PP-*ScARO7$^{FBR}$* | *ScARO7$^{FBR}$* part plasmid | This study |
| pYTK-PP-*BvCYP76AD5* | *BvCYP76AD5* part plasmid | This study |
| pYTK-PP-*PpDODC* | *PpDODC* part plasmid | This study |
| pYTK-PP-*ScTYR1* | *ScTYR1* part plasmid | This study |
| pYTK-PP-*PpTA* | *PpTA* part plasmid | This study |
| pYTK-PP-*DpTA* | *DpTA* part plasmid | This study |
| pYTK-PP-*ZmTA* | *ZmTA* part plasmid | This study |
| pYTK096 | *URA3* integration vector | [13] |

**Supplementary Table 5 – *S. cerevisiae* integration sites utilized in this study**

| Target site ID | Target site sequence[a] | Reference |
|---|---|---|
| 106a | ATACGGTCAGGGTAGCGCCCTGG | [14] |
| 308a | CACTTGTCAAACAGAATATAAGG | [14] |
| 911b | GTAATATTGTCTTGTTTCCCTGG | [14] |
| *ADH6* | AAAAGCACCAACAGTTCTCGAGG | This study |
| *ARI1* | AAGTTGCATAGAATAAATTCCGG | This study |
| *GCY1* | GAAAGCTCGGTTTCTACTCTAGG | This study |
| *GRE2* | CTTAGTTTCCCAACTATATAAGG | This study |
| *GRE3* | CCTCGAAGTTACTACTTCTAGGG | This study |
| *HFD1* | TTAGTGATTTAATTAGATGGTGG | This study |
| *YDR541C* | GATCGCATGCATGTTCGCTGCGG | This study |
| *YGL039W* | TCTCAATTGGCTATCCAAAAAGG | This study |
| *YPR1* | GAATTGAATTGCAACCAAATAGG | This study |

[a] PAMs are underlined
